# Gonococcal aggregation causes upregulation of genes involved in antibiotic tolerance

**DOI:** 10.1101/2025.01.17.633513

**Authors:** Sebastian Kraus-Römer, Isabelle Wielert, Isabel Rathmann, Thorsten E. Volkmann, Paul G. Higgins, Berenike Maier

## Abstract

Aggregation and biofilm formation can increase the tolerance of bacteria to external stressors, including antibiotic treatment. While resistant bacteria grow at an elevated drug dose, tolerant bacteria survive longer-term treatment. The mechanisms by which aggregation confers tolerance are insufficiently characterized for most organisms, including the human pathogen *Neisseria gonorrhoeae*. We hypothesize that bacterial aggregation causes upregulation of genes involved in tolerance and that deletion of these genes increases killing rates during antibiotic treatment. To test this hypothesis and identify genes involved in gonococcal tolerance, we compared the transcriptome of aggregating and planktonic *N. gonorrhoeae* strains. In general, the transcriptome analysis shows that aggregation causes a strong upregulation of prophage-related genes and a shift towards anaerobic respiration. We generated deletion strains for the twenty most upregulated genes and measured their killing kinetics during treatment with the clinically relevant antibiotics ceftriaxone or ciprofloxacin. We identified five genes and one multigene segment that are involved in gonococcal antibiotic tolerance. These include prophage genes whose deletion affects tolerance differently in aggregating and planktonic strains. Furthermore, deletion of genes encoding a putative multi-drug efflux pump, an alcohol dehydrogenase, and a DNA repair protein reduces tolerance. In summary, we have identified multiple genes that affect antibiotic tolerance and are upregulated in response to aggregation.

**Author summary:** Often bacterial infections recur after antibiotic treatment because not all of the bacteria were killed. The ability to survive treatment by bactericidal drugs is termed tolerance. It is well established that aggregation can increase tolerance by reducing growth and metabolism. However, the genes involved in tolerance are not well characterized, especially in the human pathogen *Neisseria gonorrhoeae*. Here, we aim to identify such genes by following the hypothesis that aggregation upregulates genes that cross-protect *N. gonorrhoeae* from antibiotic treatment. We show that prophage-associated genes are strongly upregulated in aggregates and that deletion of various phage genes affects tolerance to the currently administered drug, ceftriaxone. We identify three additional genes belonging to different functional classes whose deletion reduces tolerance to ciprofloxacin. Our study is an important step towards understanding the molecular mechanisms of gonococcal antibiotic tolerance. In particular, we propose that prophages could serve as a target for the treatment of tolerant gonococcal infections.

## Introduction

While antibiotic resistance has been intensively studied, antibiotic tolerance has emerged as a serious yet less studied factor contributing to treatment failures (1). When bacteria are intermittently treated with bactericidal antibiotics, often a small fraction of the population will survive the treatment. Since killing is a stochastic process that occurs at a specific rate, the fraction of survivors follows characteristic killing kinetics. The shallower the killing kinetics the more tolerant are the cells. Alternatively, a subpopulation of cells can enter a different physiological state, the persister state, which has a lower killing rate. Increased tolerance and persistence can evolve during repeated intermittent antibiotic treatment (2–4). Antibiotic tolerance is problematic for the treatment of infections because the surviving subpopulation can cause recurrent infections (1) and tolerance often precedes and allows for the development of antibiotic resistance (5). One of the factors influencing the killing rate is cell aggregation and subsequent biofilm formation (6, 7). Here, we investigate whether genes upregulated due to aggregation are involved in the antibiotic tolerance of *Neisseria gonorrhoeae*, the causative agent of gonorrhea, one of the most common sexually transmitted infections worldwide (8).

The development of antibiotic resistance in *N. gonorrhoeae* is well documented (9, 10) and many genetic determinants underlying resistance are known (11, 12). In contrast, little is known about antibiotic tolerance of *N. gonorrhoeae.* Recently, tolerant strains of *N. gonorrhoeae* have been isolated and detected from patients and in WHO reference strains (13, 14). In several studies, tolerance to antibiotics has been investigated in vitro (15–17), but the molecular mechanisms and genes involved in tolerance remain largely elusive (18).

In other bacterial species, several genes have been identified that contribute to antibiotic tolerance (1), most of which are related to slow growth and reduced metabolism. The corresponding metabolic pathways include purine synthesis, stringent response, the TCA cycle, as well as the electron transport chain, but we assume that these represent only a fraction of genes and molecular mechanisms that mediate tolerance (1). Slow growth and reduced metabolism are pronounced in bacterial biofilms and are therefore likely to contribute to the protective effect of biofilm formation (6, 7). Moreover, there is evidence that aggregation triggers stress responses that cross-protect the bacteria from antibiotic treatment, including the stringent response and SOS response (19, 20).

*N. gonorrhoeae* (gonococci) form biofilms in vitro (21) and there is evidence for in vivo formation of biofilms (22). In vitro and on epithelial cell layers, *N. gonorrhoeae* quickly forms spherical aggregates, also known as microcolonies (23–25). The aggregation process is mediated by retractile type 4 pili (T4P) (26), which are present in large numbers at the gonococcal cell surface (27). These dynamic polymers create an attractive force between neighbouring cells (28–30) and govern the mechanical properties of the aggregates (28, 31–33). We have shown previously that within hours, cells within a microcolony differentiate into two subpopulations: a proliferating and highly energized subpopulation at the periphery of the microcolony, and a slowly growing subpopulation with reduced membrane potential at the centre (17, 34). These subpopulations exhibit different levels of tolerance to antibiotics (16, 17). It is currently unclear how gene expression changes during the early stages of gonococcal biofilm development. However, previous studies have characterized longer-term differences in gene expression between planktonic and biofilm-attached cells of a clinical *N. gonorrhoeae* isolate. Cells were grown in a flow chamber for two days and transcription in adherent cells and cells of the supernatant were compared using RNA microarray analysis (35). Under these experimental conditions, 3.8 % of the genome was differentially expressed. Genes encoding enzymes belonging to the anaerobic respiratory metabolism (*aniA*, *norB*, *ccp*) were among those that were strongly upregulated, while genes of the *nuo* cluster involved in aerobic respiration were among those that were downregulated. Deletion of genes involved in anaerobic respiration attenuated biofilm formation, suggesting that gonococci assume an anaerobic metabolism within the biofilm (35). Proteins belonging to the nitrite reduction pathway were also detected as upregulated in a proteomics study (36). Apart from this class of proteins / genes, there was little overlap between the biofilm transcriptome and the proteome although the experimental setup for biofilm formation was similar between both studies. The proteome study showed additional differential regulation of proteins involved in energy metabolism, protein synthesis, and cell envelope proteins (36). In the experimental setup used in these studies, the planktonic cells were sampled from the supernatant. Therefore, it is unclear whether they had recently dispersed from the biofilm and how the prehistory of biofilm growth and subsequent release affects gene expression.

Aggregation enhances antibiotic tolerance of *N. gonorrhoeae* to ceftriaxone (15, 37) and ciprofloxacin (37). It is unclear, however, which genes are involved in tolerance and which of them are related to aggregation. Here, we aim at identifying genes that affect gonococcal tolerance. It has been shown for other bacterial species that differential gene regulation triggered by aggregation can provide cross-protection against antibiotic treatment (19, 20). With this knowledge in mind, we set out to investigate the effect of aggregation-induced upregulation on the antibiotic tolerance of *N. gonorrhoeae*. In the first step, we examined which genes are differentially transcribed between planktonic and aggregating cells of *N. gonorrhoeae* and identify the twenty most upregulated genes. We then investigate the effects of deletion of these genes on tolerance to ceftriaxone and ciprofloxacin. To find out whether these effects require aggregation, we compare the effects in the background of aggregating and planktonic cells. We show that deletion of various genes associated with prophages impact tolerance differently in aggregating and planktonic strains. In aggregating strains, genes involved in drug efflux, DNA repair, and metabolism affect tolerance. The identification of these genetic determinants of tolerance will help to build a picture of the molecular mechanisms of antibiotic tolerance in *N. gonorrhoeae*.

## Results

### Aggregation causes profound changes in transcription

T4P can attractively interact and by this govern gonococcal aggregation (23, 28, 30). Here, we investigate how aggregation affects transcription by RNA sequencing of aggregating and planktonic strains. The two aggregating strains used in this study contain different variants of the gene encoding the major pilin *pilE*, wt_agg_ and wt_agg2_ (37) (S1 Table). Two of the planktonic strains (wt_plank,_ wt_plank2_) generate type 4 pili which do not mediate aggregation, but are otherwise functional (37). Furthermore, we used a planktonic strain that lacks *pilE* and cannot form pili (*ΔpilE*). For transcriptome analysis, we allowed the gonococcal strains to grow for 10 h. wt_agg_ and wt_agg2_ formed colonies at this time point, while wt_plank,_ wt_plank2_, and the *ΔpilE* strains are planktonic (37). For all strains, RNA was isolated and sequenced.

In the first step, we evaluated clustering of the RNA-seq reads using principal component analysis (PCA). We found that the two aggregating strains and the three planktonic strains cluster separately in the first principal component (PC1) (Fig. 1A), which encompasses 78 % of the total variance. This result indicates that aggregation strongly affects transcription of gonococci. Interestingly, wt_plank2_ replicates cluster separately from all the other strains along the second principal component (PC2), suggesting that the wt_plank2_ expression data across all genes differs significantly from the other samples. The genes that have the largest influence when calculating the principal components in the new basis include phage-associated genes (NGFG_01274, NGFG_01290), an ATP-dependent RNA helicase (NGFG_01150) and two opacity proteins (NGFG_02259, NGFG_02351). When the genes with the highest effect on the PCA are excluded from the analysis, the clustering of conditions remains stable and similar to Fig. 1A, indicating that the clustering is caused by global strain differences and not by individual genes.

**Fig. 1.**
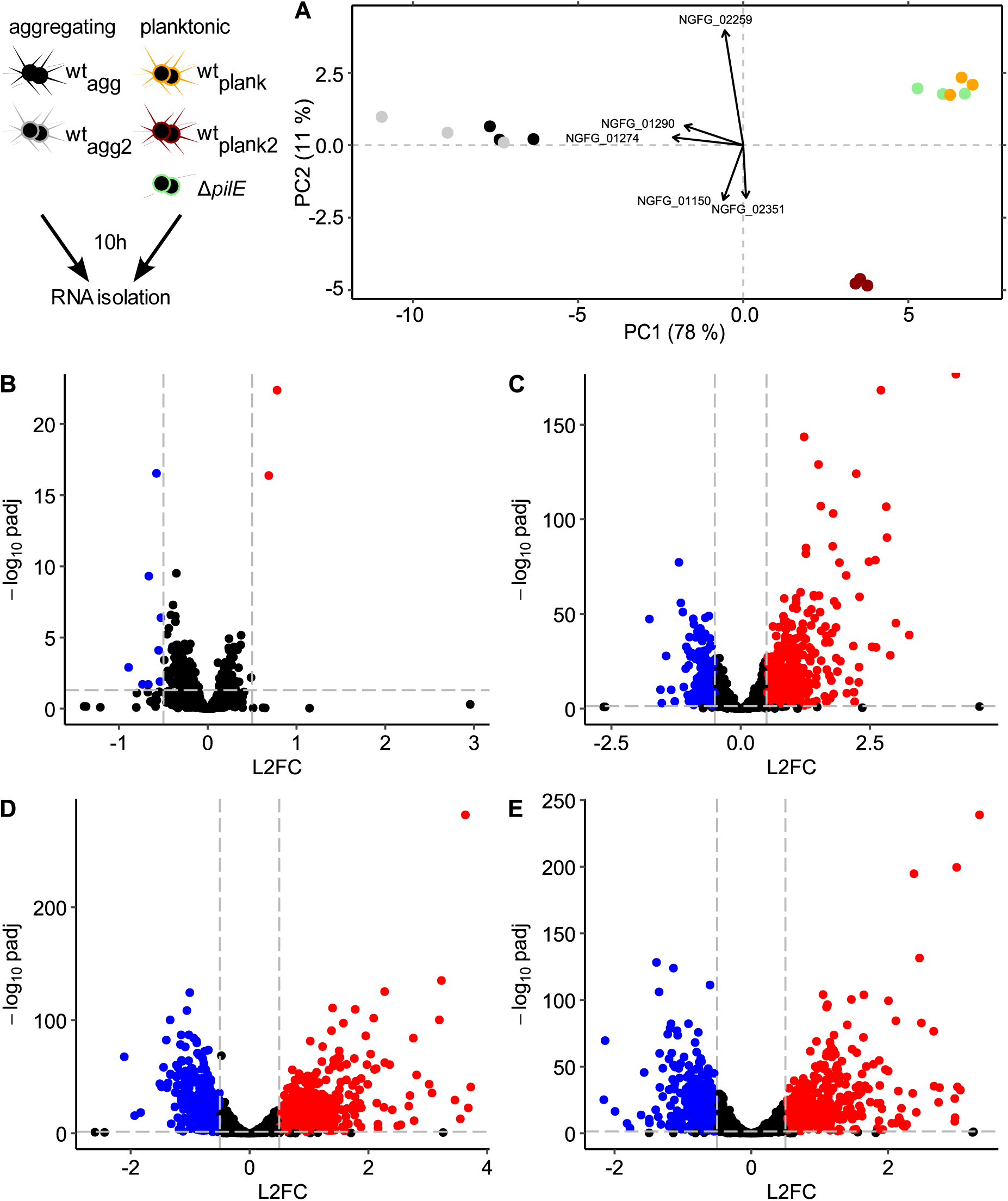
Overview of the transcriptomes of aggregating and planktonic strains. A) PCA analysis. wt_agg_ (black), wt_agg2_ (gray), wt_plank_ (orange), wt_plank2_ (red), and *ΔpilE* (green). Volcano-plots showing the adjusted p-value as a function of the log_2_-fold change (L2FC) for wt_agg_ versus B) wt_agg2_, C) wt_plank2_, D) wt_plank_, E) *ΔpilE*. Dashed lines depict the cut-offs for significant differential transcription. Blue: significantly down-regulated genes, red: significantly up-regulated genes.

In the next step, we compare the transcription of the aggregating wt_agg_ strain with each of the four T4P-replacement strains. We consider genes to be differentially regulated if − 0.5 ≤ *log*_2_ *fold change* ≤ 0.5 and the adjusted p-value is *p* < 0.05. Between the two aggregating strains wt_agg2_ and wt_agg_, we find small differences in their transcriptome (Fig. 1B) as only 0.1 % of all genes are upregulated and 0.4 % are downregulated significantly. Comparing wt_agg_ to the planktonic strains reveals considerable differences in expression, with between 16 % and 2.7 % of genes being upregulated and between 9.7 % and 18.9 % of genes being downregulated (Fig. 1C-E).

Overall, RNA-seq analysis shows strong differences in gene expression between the aggregating and planktonic strains.

### Transcription reveals upregulation of prophage-related genes and indicates a shift to anaerobic respiration

We assessed whether genes associated with specific biological functions were collectively upregulated or downregulated between the aggregating and planktonic strains. To this end, we performed a functional enrichment analysis for each strain using KEGG categories, as described previously (27, 38) (see Methods for details) (Fig. 2, S1 Fig.).

**Fig. 2.**
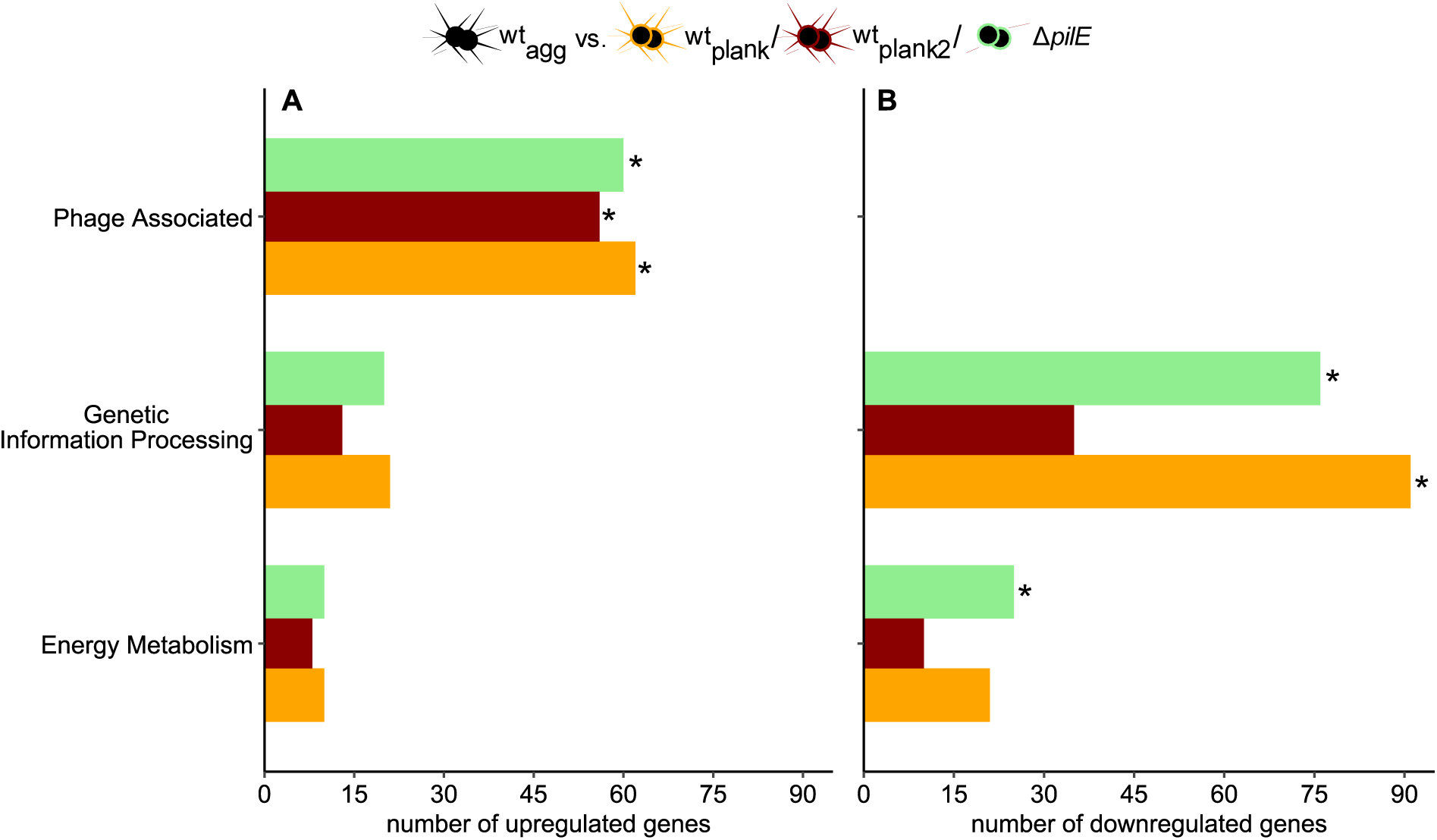
Functional enrichment of up- and down-regulated genes in the wt_agg_ strain versus the *ΔpilE* (green), wt_plank2_ (red), and planktonic strains wt_plank_ (orange). Here, only significant KEGG categories are shown and the number of up- and down-regulated genes is computed and depicted as bar plots. Categories are analysed for enrichment with a Fisher’s exact test and multiple testing correction is performed with the Bonferroni method. Significantly enriched categories are highlighted with * (p-value ≤ 0.01).

We find that the category of phage-associated genes is enriched among the upregulated genes when comparing wt_agg_ to all three planktonic strains (Fig. 2). The *N. gonorrhoeae* genome contains four tailed double-stranded prophages (39) (S2 Fig. A). In wt_agg_, genes belonging to all of these prophages are significantly upregulated when compared to the planktonic strains (S2 Fig. B-E), indicating that aggregation causes upregulation of genes within the double-stranded prophages. Upregulation is most prominent for genes that belong to prophages Φ1 and Φ3.

The categories of genetic information processing and energy metabolism are enriched among the downregulated genes when comparing wt_agg_ to some of the planktonic strains (Fig. 2). Among the genes belonging to the energy metabolism category, genes involved in denitrification (*aniA* (NGFG_02154), *norB* (NGFG_02153), and *nos* genes) are upregulated, while genes involved in aerobic respiration (*nuo* genes) tend to be down-regulated. This indicates a shift from aerobic to anaerobic respiration in agreement with a previous report (35). For the category of genetic information processing, genes encoding ribosomal genes are among the most strongly downregulated genes within the aggregating strains. These genes include *rpl* and *rpm* genes encoding for the 50S subunit as well as *rps* genes encoding for the 30S subunit.

We define the aggregation transcriptome as the intersection of genes that are differentially regulated in all three planktonic strains compared to strain wt_agg_. Within this intersection, 239 genes are consistently upregulated and 138 genes are consistently downregulated (Fig. 3). In terms of functional KEGG categories, only the category of phage-related genes is significantly overrepresented.

**Fig. 3.**
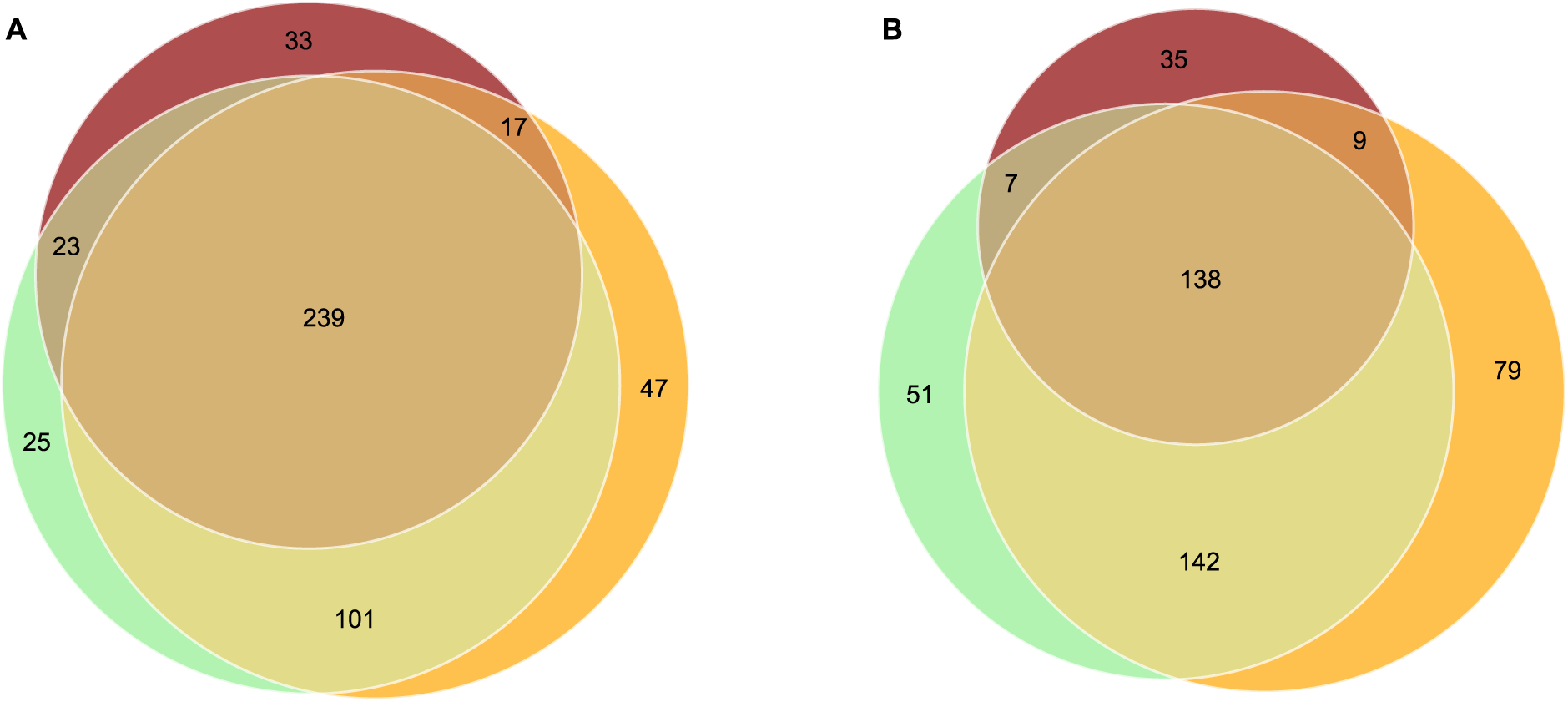
Intersection of genes that are differentially transcribed by colony-forming strains compared to planktonic strains. A) upregulated genes, B) downregulated genes. wt_agg_ vs. wt_plank_ (orange), wt_plank2_ (red), and *ΔpilE* (green).

We conclude that 239 genes are upregulated and 138 genes are downregulated due to aggregation. The upregulated genes are functionally enriched in phage-associated genes.

### Deletion of various genes upregulated in aggregates affects antibiotic tolerance to ciprofloxacin

To better understand the mechanisms causing increased tolerance, we scrutinized the effect of deleting the twenty most strongly upregulated genes of the aggregation transcriptome (Fig. 3) (S1 Table). First, we generated deletion strains within the wt_agg_ background. When consecutive genes on the genome belonged to these highly upregulated genes, they were deleted together in one strain. We have previously shown that the wt_agg_ strain multiplies nearly exponentially during the initial 10 h (37). To find out whether the gene deletions have a strong effect on the growth dynamics, we compared the number of cells after 10 h of growth between the different mutants. None of the deletion strains showed a significant growth defect (S3 Fig.). Furthermore, we assessed the minimal inhibitory concentrations of ciprofloxacin and ceftriaxone and found no significant changes except the *ΔrecN* strain that showed a two-fold reduction of the ciprofloxacin MIC (S2 Table).

For each deletion strain, we characterized the killing kinetics at its 600-fold MIC. This concentration was chosen for consistency with an earlier work in which we found strong effects of aggregation on the killing kinetics (37). By comparing killing kinetics over a time range of 9 h between the deletion strains and their parental strains, we determined deletions that altered the killing kinetics. We calculated the fraction of surviving cells (f) by dividing the CFU at the start of the antibiotic treatment by the CFU after treatment for either 3, 6 or 9 h (*f* = *CFU*_*start*_/*CFU*_*treatment*_) (S4 Fig.). Each condition was characterized in at least three independent experiments and we evaluated the significance by weighting with the number of replicates (40).

We discovered three gene deletions that significantly accelerated killing of wt_agg_ by ciprofloxacin (Fig. 4A). These deletions include the DNA repair gene *recN* (NGFG_00464), the gene encoding the putative *drug:H^+^* antiporter (NGFG_00826), and the alcohol dehydrogenase gene *adh* (NGFG_01080) (Fig. 4A). After the entire 9-hour treatment period, the deletion of each of the three genes reduces the fraction of surviving cells by a factor of 5 to 10.

**Fig. 4.**
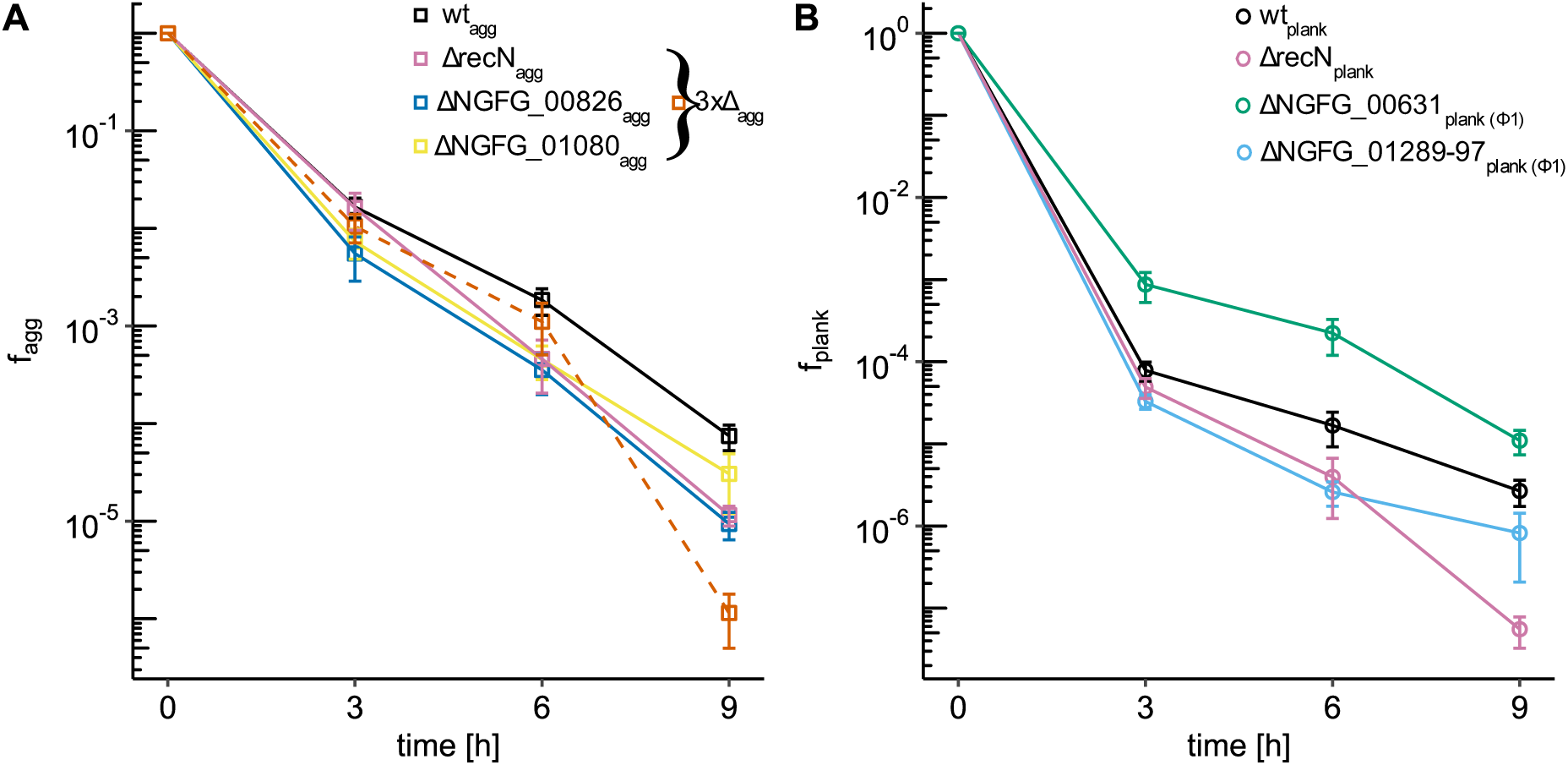
Killing kinetics of deletion strains in aggregating (A, f_agg_) and planktonic (B, f_plank_) background under treatment with 2.4 µg/ml ciprofloxacin. Only strains for which the killing kinetics are significantly different from the kinetics of the parental strain are shown. Shown is the fraction of surviving cells *f = CFU_start_/CFU_treatment_*. A) Fractions of surviving aggregating cells (squares) with gene deletions and their respective parental strain. wt_agg_ (Ng150, black), *ΔNGFG_01080*_agg_ (Ng281, yellow), *ΔrecN*_agg_ (Ng261, pink), *ΔNGFG_00826*_agg_ (Ng262, blue), *3xΔ*_agg_ (pooled data of Ng321 and Ng325, dashed orange). B) Fractions of surviving planktonic cells (circles) with gene deletions and their respective parental strain. wt_plank_ (Ng240, black), *ΔrecN*_plank_ (Ng275, pink), *ΔNGFG_00631*_plank_ (Ng288, green), *ΔNGFG_01289-97*_plank_ (Ng300, cyan).

We wondered whether a combinatorial deletion of genes significantly affecting the survival during ciprofloxacin treatment would have more severe effects. Therefore, we constructed a triple deletion strain *ΔrecN ΔNGFG_01080 ΔNGFG_00826_agg_* (3xΔ_agg_). Compared to the single deletion strains (Fig. 4A), the triple deletion strain did not show accelerated killing (Fig. 4A). However, the number of cells was significantly reduced after 10 h of growth in that strain (Fig. S9A), suggesting that the cells grew more slowly. Since slow growth is often associated with increased tolerance (41), we cannot rule out that faster killing caused by the gene deletions and slower killing caused by reduced growth cancel each other out.

Next, we addressed the question of which gene deletions affect tolerance in the planktonic strains. We deleted the same 20 genes (S1 Table) in the planktonic background (wt_plank_) and determined the killing kinetics of the deletion strains (S5 Fig.). Most deletions showed no strong growth effect after 10 h with the exceptions of strains *ΔNGFG_00631*_plank_ and *ΔNGFG_01491*_plank_ that had a slightly but significantly higher cell number at that growth stage (S3 Fig.). Again, we found that deletion of *recN* significantly accelerated killing (Fig. 4B), showing that this gene is important for tolerance to ciprofloxacin both in aggregates and for planktonic strains. Furthermore, we discovered that deletion of the multigene segment NGFG_01289-97 from prophage Φ1 accelerated the killing of planktonic strains. Interestingly, deletion of the phage related gene NGFG_00631 (Φ1) significantly slowed killing of the planktonic strain.

We conclude that deletion of genes encoding for a putative drug antiporter, a DNA repair protein, and an alcohol dehydrogenase reduce tolerance against ciprofloxacin.

### Phage-related genes affect tolerance towards ceftriaxone

Next, we addressed the question of whether deletion of the 20 most upregulated genes of the aggregation transcriptome affects killing during the treatment with a different class of antibiotics. We treated the deletion strains with ceftriaxone at 600-fold MIC (4.8 µg/ml). Under ceftriaxone treatment, we did not identify deletions that significantly enhanced killing in the aggregating background (S6 Fig.). However, deletion of the phage-related genes NGFG_01289-97 significantly increased the fraction of surviving cells (Fig. 5A). Interestingly, the same deletion reduced the fraction of surviving bacteria in the planktonic background (Fig. 5B, S7 Fig.). This shows that the effects of these phage-related genes on ceftriaxone susceptibility depend on aggregation. Additionally, deletion of a putative terminase gene of prophage Φ1 (NGFG_01302) significantly reduced survival. Finally, deletion of the phage related gene NGFG_00631 (Φ1) significantly slowed killing under ceftriaxone treatment.

**Fig. 5.**
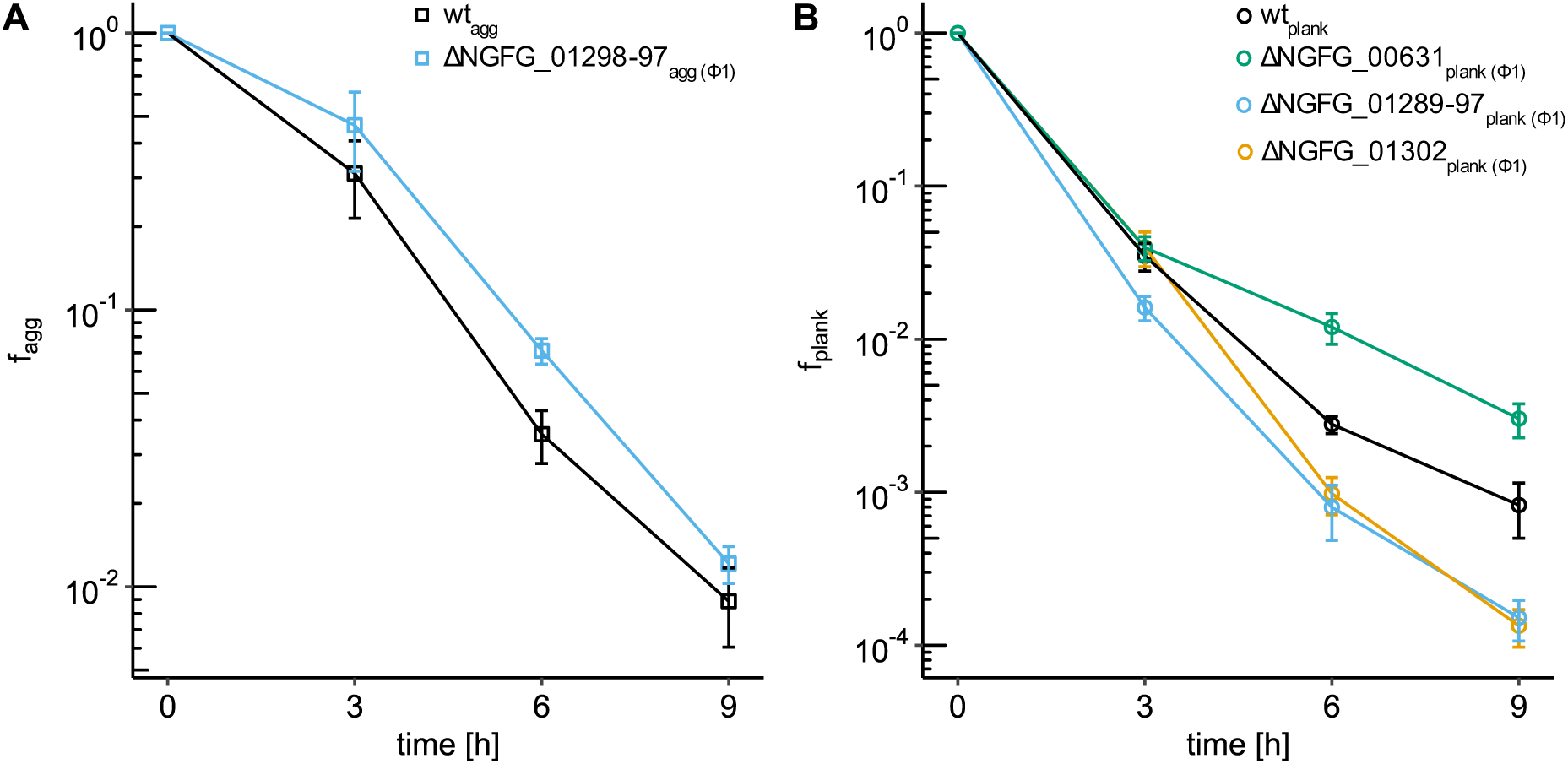
Killing kinetics of deletion strains under treatment with 4.8 µg/ml ceftriaxone. Only strains whose killing kinetics is significantly different from the kinetics of the parental strain are shown. Shown are the fraction of surviving cells *f = CFU_start_/CFU_treatment_*. A) Fractions of surviving aggregating cells (squares) with gene deletions and their respective parental strain. wt_agg_ (Ng150, black), *ΔNGFG_01289-97*_agg_ (Ng273, cyan). B) Fractions of surviving planktonic cells (circles) with gene deletions and their respective parental strain. wt_plank_ (Ng240, black), *ΔNGFG_01302*_plank_ (Ng302, orange), *ΔNGFG_00631*_plank_ (Ng288, green), *ΔNGFG_01289-97*_plank_ (Ng300, cyan).

We conclude that the deletion of genes related to prophage Φ1 impacts the tolerance to ceftriaxone. In particular, deletion of a multigene segment affects the tolerance of aggregating and planktonic strains in opposite ways highlighting its lifestyle-specific effect.

## Discussion

In this study, we used transcriptomic data to identify genes involved in antimicrobial tolerance. We found five genes and one multigene segment that modify tolerance to ceftriaxone and/or ciprofloxacin. Genes influencing ciprofloxacin tolerance have different putative functions, whereas genes affecting ceftriaxone tolerance are associated with one of the prophages on the gonococcal genome. Our results point to tolerance mechanisms that differ from the reduction in cell growth and metabolism that has been demonstrated as a common theme in many studies as the cause of antibiotic tolerance (1, 7).

### Differential expression in aggregates indicates anaerobic lifestyle

The transcriptome of the colony-forming isogenic strains differs significantly from that of the planktonic strains. The only genetic difference between the strains was the pilin variant that controls the attraction between cells (37). Thus, our results show that pilin antigenic variation strongly affects the transcriptional pattern of gonococci by regulating aggregation.

We found that genes involved in aerobic respiration showed a tendency to be downregulated in aggregating strains while genes involved in anaerobic respiration were more likely to be upregulated. Therefore, we compared our data with a study reporting on the anaerobic stimulon of *N. gonorrhoeae* F62 (42) and found a considerable overlap. Of the 196 differentially expressed genes during anaerobic conditions, we found 169 annotated orthologues in MS11. Up to 41 % of those genes were similarly regulated under our conditions. The gene categories that contain the most similar regulated genes belonged to regulation, phages, small molecules, and macromolecular biosynthesis. Genes that were strongly upregulated under both conditions include *aniA*, *nosR*, *recN*, *NGFG_01491*, and several genes encoding prophages Ngo Φ1-4. The observation that prophage genes were strongly upregulated under both conditions suggests a role for prophages in gonococcal biofilm development. There are interesting differences between the transcriptional responses to aggregation and oxygen limitation. For example, we show that various genes involved in iron and sulfate acquisition are upregulated in aggregates, while these genes tend to be downregulated during purely anaerobic conditions (42).

We compared our results with earlier works on gene expression in gonococcal biofilms, which used microarray and proteomic methods (35, 36). Those studies focused on mature biofilms and used a different method for comparing planktonic and biofilm-forming bacteria. Their studies compared cells living in biofilms to cells that had dispersed from the biofilm. PCA analysis of *P. aeruginosa* biofilms showed that dispersed cells cluster separately from the biofilm and planktonic cells, suggesting that dispersed cells represent a distinct stage in the transition from the biofilm to the planktonic lifestyle (43). Despite the different methods used in both studies (35, 36), their results were similar to our findings in terms of differential expression of genes involved in anaerobic respiration and the downregulation of genes involved in protein synthesis and energy metabolism. This shows that this transcriptional response to oxygen limitation as a consequence of aggregation is robust across experimental conditions of biofilm formation.

### Genes with diverse functions affect tolerance of aggregating gonococci to ciprofloxacin

In this study, we started from the assumption that during aggregation, specific genes that also protect bacteria from antibiotic treatment are upregulated. Under ciprofloxacin treatment, we identified three genes from different functional classes whose deletion affected tolerance of gonococcal aggregates.

Deletion of *NGFG_00826* in the aggregating strain resulted in a less ciprofloxacin-tolerant phenotype. *NGFG_00826,* is annotated as the major facilitator superfamily (MFS) “drug:H+ antiporter”. The MFS transporters are secondary active transporters with a broad spectrum of substrates (44). In bacteria, these systems are used for nutrient uptake but also for export of toxic compounds such as antibiotics or heavy metals (45). MFS transporters contribute to antibiotic resistance in *E. coli* and *S. aureus* (46, 47). It is likely that this MFS transporter is also involved in extrusion of ciprofloxacin in *N. gonorrhoeae*. Similarly, an ABC transporter was described in *P. aeruginosa* that was upregulated in biofilms and increased tolerance to antibiotics, including ciprofloxacin (48). We propose that *NGFG_00826* drug transporter is upregulated in response to aggregation and cross-protects gonococci from ciprofloxacin treatment.

Moreover, deletion of a gene encoding an alcohol dehydrogenase (ADH), *NGFG_01080,* resulted in a less ciprofloxacin-tolerant phenotype in the aggregating strain. ADH are enzymes that convert alcohols to aldehydes and *vice versa*. Various studies showed that downregulating metabolic genes can increase tolerance (1). Here we observe the opposite: *adh*, a gene involved in carbon metabolism, is upregulated and its deletion reduces tolerance. ADH may be involved in maintaining redox balance and detoxifying reactive oxygen species, which often result from antibiotic treatment (49). Reduction equivalents such as NAD^+^/NADH are converted during the oxidation of alcohols. Through the deletion of *NGFG_01080,* we might disturb the redox balance within the cells, thereby generating aggregates that are less resilient to antibiotic stress and reducing *N. gonorrhoeae*’s ability to deal with reactive oxygen species. It is tempting to speculate that *adh* is upregulated during aggregation to detoxify reactive oxygen species, and that this effect also protects the bacteria from antibiotic treatment.

The third gene whose deletion decreased tolerance to ciprofloxacin was *NGFG_00464,* which encodes the DNA repair protein RecN. Its deletion reduced tolerance in both aggregating and planktonic backgrounds even though the expression levels of *recN* were significantly lower in the planktonic strains. Since ciprofloxacin causes DNA damage, it was expected that deletion of a DNA repair gene would affect survival and growth under ciprofloxacin treatment.

In summary, we have identified three genes that influence the tolerance of aggregating gonococci to ciprofloxacin. Since these genes are among the most upregulated genes in aggregates compared to planktonic bacteria, it is reasonable to assume that their upregulation contributes to the protective effect of aggregation.

### Prophage **Φ**1 genes influence tolerance in aggregating and planktonic bacteria

In our study, predominantly prophage-encoded genes were upregulated in the aggregated state. Recently, it was shown that prophage induction in other biofilm-forming species releases matrix components (50, 51). Activation of prophages may contribute to the stability of the aggregates by actively integrating phages into the matrix, as is the case in other pathogenic species (52, 53). We find that none of the gene deletions in the wt_agg_ background affected its ability to form aggregates in liquid culture (S10 Fig.), suggesting that a different mechanism is at play in our system. Very little is known about the role of prophage activation of *N. gonorrhoeae*. In an early genomic study (54), five regions of strain FA1090 were assigned to dsDNA prophages Φ1-Φ5 and DNA from Φ1 was detected, suggesting that this phage is active. The latter region is also present in the derivatives of strain MS11 studied here. We used PHASTEST (39) and predicted four prophages on the genome (S2 Fig. A). The region corresponding to Φ1 is predicted by the software to be intact. All the genes we have identified as influencing tolerance belong to this prophage and many of them are upregulated in aggregates (S2 Fig. B). However, we show that in the planktonic strains, deletion of genes belonging to this prophage affects tolerance. The reason for this could be that some genes located in the prophage region affect bacterial physiology and thus tolerance. As discussed below, we consider it is more likely that the phage can be activated in the planktonic state, possibly by antibiotic treatment.

Deletion of *NGFG_00631* increased the tolerance of planktonic strains to ceftriaxone and ciprofloxacin. BLAST analysis of *NGFG_00631* revealed high sequence identity with *cro*, a transcriptional regulator of the Cro/CI family. It has been shown that an λcro^−^ mutant cannot induce the lytic cycle of phage λ (55). The adjacent gene *NGFG_00630,* which is expressed in the opposite direction (but is not significantly upregulated or downregulated in our transcriptional analysis), shows high similarity to the gene encoding the repressor protein CI which is crucial for maintaining the lysogenic cycle of phage λ. We hypothesize that these proteins resemble the two regulatory proteins responsible for the bistable switch of temperate bacteriophages. Assuming that phage Φ1 can actively enter the lytic cycle, deletion of *NGFG_00631* would prevent the transition to the lytic cycle and explain the enhanced survival of planktonic mutants during antibiotic treatment. This is consistent with the increased cell number after ten hours of growth (S3 Fig.). However, we do not observe an enhanced survival of the corresponding aggregating *NGFG_00631* deletion strain. We hypothesize that lytic cycle-induced cell lysis provides a source of extracellular DNA that can be integrated into the extracellular matrix (22, 56). Thus, lysis in a subpopulation of gonococcal aggregates could enhance the structural integrity of aggregates and subsequently impact survival during antibiotic treatment.

The deletion of the genes *NGFG_01289-97* showed the most ubiquitous effect on gonococcal tolerance. In the planktonic strain, this deletion led to reduced tolerance to ciprofloxacin and ceftriaxone. When the same stretch of genes was deleted in the aggregating strain, the mutants showed a higher tolerance to ceftriaxone than the wt_agg_. According to BLAST analyses, these genes encode a variety of phage tail and connector proteins. These proteins are essential for phage assembly (57), but their deletion may not prevent cell lysis once the lytic cycle has been activated. Lack of these components could lead to conditions similar to those of an abortive infection during a phage infection, in which the cell harbouring the unassembled phage dies before phage maturation (58). This effect could prove detrimental to planktonic cells, especially under antibiotic stress. However, the aggregated counterpart might benefit from cell lysis prior to phage assembly, as more cell debris could be integrated into the extracellular matrix, especially free unpackaged DNA. Deletion of *NGFG_01302* may have similar effects, as it encodes the small terminase subunit TerS. The absence of this essential subunit for DNA recognition and the initiation of the packaging process could interfere with virus assembly, which would have the same effect as deletion of *NGFG_01289-97*.

We conclude that multiple genes related to prophages influence antibiotic tolerance. Importantly, deletion of a stretch of genes which is most likely essential for phage assembly leads to opposite results in aggregating and planktonic strains when treated with ceftriaxone. This suggests that the role of prophage genes in tolerance depends on bacterial aggregation.

## Conclusion

Gonococcal aggregation increases antibiotic tolerance (15, 37). Therefore, genes that affect aggregation, including *pilE*, *pptA*, *pilT*, *opa*, and *lgtE*, modulate tolerance (15, 16, 37). Most mechanisms that cause bacterial tolerance are based on reducing growth rates or the slowing down of metabolic pathways (1, 7). We have previously shown that aggregation decelerates the growth rate of gonococci (37), and it is conceivable that growth rate reduction directly increases tolerance. Also, consistent with a slowed metabolism, there is evidence that point mutations in the gene encoding phosphopyruvate hydrase *eno* may be involved in tolerance (18). In this study, we identify multiple genes that affect antibiotic tolerance but not growth, of gonococci. Three genes with putative functions in drug efflux, DNA repair, and carbon metabolism and/or ROS detoxification are strongly upregulated in aggregates and involved in increasing the tolerance of those aggregates to ciprofloxacin. In addition, two genes and one multigene segment related to prophage Φ1 affect tolerance to ceftriaxone and ciprofloxacin. Although gonococcal phages are poorly characterized (54), our results are most consistent with the interpretation that phage induced lysis is detrimental to planktonic cells and, simultaneously reduces antibiotic tolerance. In contrast, our data suggest that prophages could be activated by aggregation and aggregated gonococci are less sensitive to phage activation. Overall, the mechanisms of gonococcal antibiotic tolerance are diverse and we propose that multiple mechanisms act together in gonococcal aggregates and biofilms. It will be particularly interesting to study the interplay between aggregation, prophage activation, and antibiotic tolerance in future studies.

## Materials and Methods

### Growth conditions

*N. gonorrhoeae* was grown at 37 °C and 5 % CO_2_. Gonococcal base agar was made from 10 g/l dehydrated agar (BD Biosciences, Bedford, MA), 5 g/l NaCl (Roth, Darmstadt, Germany), 4 g/l K_2_HPO_4_ (Roth), 1 g/l KH_2_PO_4_ (Roth), 15 g/l Proteose Peptone No. 3 (BD Biosciences), 0.5 g/l soluble starch (Sigma-Aldrich, St. Louis, MO), and supplemented with 1% IsoVitaleX (IVX): 1 g/l D-glucose (Roth), 0.1 g/l L-glutamine (Roth), 0.289 g/l L-cysteine-HCL x H_2_O (Roth), 1 mg/l thiamine pyrophosphate (Sigma-Aldrich), 0.2 mg/l Fe(NO_3_)_3_ (Sigma-Aldrich), 0.03 mg/l thiamine HCl (Roth), 0.13 mg/l 4-aminobenzoic acid (Sigma-Aldrich), 2.5 mg/l β-nicotinamide adenine dinucleotide (Roth), and 0.1 mg/l vitamin B12 (Sigma-Aldrich). Liquid GC media is identical to the base agar composition but lacks agar and starch.

### Antibiotics

Antibiotics were purchased from Roth, Merck, Thermo Fisher, or Hello Bio. Stock solutions of antibiotics were prepared as follows. Ceftriaxone was dissolved in DMSO and adjusted to a final concentration of 1 mg/ml. Ciprofloxacin was dissolved in 0.1 M HCl (10 mg/ml). Erythromycin was dissolved in ethanol (10 mg/ml). Streptomycin (100 mg/ml) and kanamycin (50 mg/ml) were dissolved in milliQ water. For selective plates, the final concentration of erythromycin, streptomycin and kanamycin were 5-10 µg/ml, 100 µg/ml and 50 µg/ml, respectively.

### Bacterial strains

All strains used in this study (S3 Table) were derived from the *opa-* selected VD300 strain (59). To suppress antigenic variation, the G4 motif upstream of *pilE* was deleted in all strains (60). The *pilE* variant strains from Wielert et al (37) were used and named according to their planktonic (_plank_) or aggregative (_agg_) phenotype.

### Construction of single knock-out strains

We generated deletion strains of the twenty most upregulated genes of the aggregation transcriptome (S1 Table) within the wt_agg_ background. When consecutive genes on the genome belonged to these highly upregulated genes, they were deleted together in one strain. The deletion strains were generated by replacing the gene of interest by a kanamycin resistance cassette including a promoter. Therefore, all genes residing downstream of the gene of interest within the same operon are affected by the deletion. In general, we expected genes in the same operon to have a comparable differential regulation and in such cases deleted the entire operon. In addition, we used the transcriptome study by Remmele et al. (61) to assess potential downstream effects of the gene deletions. In this study, the only deletion likely to affect a downstream gene is that of *ΔNGFG_00631,* which may affect another phage-related gene. Therefore, for most genes located in operons, our study cannot distinguish which specific gene of the operon is responsible for the effect.

Single genes or multiple consecutive genes were replaced by a kanamycin resistance cassette (aminoglycoside phosphotransferase AHM36341.1). The procedure for this replacement was the same for all mutants. In brief, the up- and downstream regions of the gene of interest were fused with a kanamycin resistance cassette and used to transform *N. gonorrhoeae*. By homologous recombination the gene of interest is replaced by the *kanR* construct.

Primers A & B and primers C & D were used to amplify the upstream and downstream region of the genes of interest from gDNA of strain wt_agg_ (Ng150). Primer E & F were used to amplify the kanamycin resistance cassette from genomic DNA of strain Ng052. The corresponding primer combinations are listed in S4 Table and the primer sequences are found in S6 Table. All amplified PCR products were purified with the PCR purification KIT (Qiagen) and then fused with the GXL-polymerase (TaKaRa). Then wt_agg_ (Ng150) or wt_plank_ (Ng240) were transformed with the fusion products. Selection was achieved by plating transformants on agar plates containing kanamycin. Correct replacement was checked via screening PCR with primers A & D and sequencing (A or D).

The gene deletions generated based on the twenty most upregulated genes in the aggregation transcriptome are listed in S1 Table. The deletion of genes may affect the expression of genes residing downstream of the deleted genes (62). We expect that genes residing within the same operon show similar patterns of differential gene expression within the gonococcal aggregates compared to planktonic cells. Therefore, we deleted all consecutive genes in a single strain if all of them were strongly upregulated. Nevertheless, in some strains we cannot exclude that the expression of genes downstream of the deleted gene are affected by the replacement of the gene of interest with the antibiotic resistance cassette which has its own promoter. Following the operon analysis of Remmele et al (61), we find only one gene that most likely resides within an operon with a downstream gene among the deleted genes (S1 Table).

### Construction of triple knock-out strain

Triple deletion strains (S3 Table) were generated by clean deletion of two genes in the background of deletion strains carrying the *kanR* resistance cassette as described above. The upstream and downstream regions of the gene of interest (each ∼1000 bp) were amplified from genomic DNA of wt_agg_ using primers cdA & cdB and cdC & cdD, respectively. The *ermC rpsL_s_* cassette was amplified from strain *T126C* step1 (Ng225) using primers cdE & cdF. The PCR products were purified with the PCR purification KIT (Qiagen) and subsequently fused with the GXL-polymerase (TaKaRa). The final fusion product was used for transformation. Selection was performed on plates containing 10 mg/ml erythromycin. Correct insertion was verified via screening PCR with primers cdA & cdD and sequencing (cdA or cdD). For the second step of the clean replacement, the upstream and downstream regions of the gene of interest (each ∼1000 bp) were amplified from genomic DNA of wt_agg_ using primers cdG & cdH and cdI & cdJ, respectively. Both PCR products were purified with the PCR purification KIT (Qiagen) and subsequently fused with the GXL-polymerase (TaKaRa). The fusion product was transformed and the strains were selected on 100 mg/ml streptomycin. Correct deletion was verified via screening PCR (cdG & cdJ) and sequencing (cdG or cdJ). All primer combinations and primer sequences are listed in S5 and S6 table, respectively.

### Determination of MIC

The minimal inhibitory concentrations (MIC) of the antibiotics ceftriaxone and ciprofloxacin were determined for each strain (S2 Table). 10 µl OD_600_= 0.1 cells were inoculated into 1 ml GC-medium in 48 well plates (Greiner Bio-One) and supplemented with doubling dilutions of antibiotics. Bacteria were grown in an Infinite M200 plate reader (Tecan) at 37°C, 5% CO_2_ with a shaking period of 2 min per OD measurement cycle. The lowest concentration of an antibiotic without detectable growth (OD_600_ ≤ 0.1) after 24 h was determined as the MIC of the respective antibiotic.

### Bacterial survival assay

Cultures were inoculated as described for the characterization of the MIC, and grown for 10 h in an Infinite M200 plate reader (Tecan) (37°C, 5% CO_2_ with a shaking period of 2 min per OD measurement cycle). After 10 h, we added antibiotics to each well except the control wells. We incubated the bacteria with 2.4 µg/ml ciprofloxacin or 4.8 µg/ml ceftriaxone for 0 h, 3 h, 6 h, or 9 h. Subsequently, the complete culture was transferred into a 1.5 ml reaction tube and cells were pelleted by centrifugation. The supernatant was discarded and 500 µl GC media and 500 µl milliQ water were added. The mixture was vortexed for 2 min. Before plating and each dilution step cells were vortexed rigorously to dissolve aggregates. Then serial dilutions were plated and incubated for 48 h (37°C, 5% CO_2_). To determine the fraction of surviving cells colony forming units (CFU) of the different treatment periods (3 h, 6 h, 9 h) were divided by the CFU of the control (0 h).

Statistical analysis of the killing kinetics was performed via a combined p-values method for discrete data (40). We have used the Mann-Whitney U test to compare single time-points followed by the combination of the p-values via Mudholkar and George combining method (63).

For both antibiotic treatments, we assessed whether the introduced kanamycin allele might affect survival. We found that in the planktonic background, which is more susceptible to antibiotic treatment, the introduction of the kanamycin resistance gene in the intergenic *aspC-lctP* locus did not affect the survival compared to wt_plank_ (S8 Fig. B, C). Furthermore, we show that neither the carrying capacity of lctP:kanR:aspC_plank_ nor *ΔpilE* is significantly different from wt_plank_ (S8 Fig. A)

### Aggregation assay in liquid media

To check whether mutants in the wt_agg_ background maintain the ability to form aggregates in liquid media we imaged the cultures (S10 Fig.). We adjusted the OD_600_ to 0.1 and added 100 µl of cells to 200 µl of fresh GC media in an ibidi 8-well. Cells were grown for 5 h (37°C, 5% CO_2_) prior to imaging. Images were recorded with an inverted Nikon Eclipse Ti with an ORCA camera model (40x magnification).

### RNA isolation and sequencing

To compare the transcriptional changes between the planktonic and the biofilm lifestyle of *N. gonorrhoeae*, different strains were grown in 1 ml GC medium in a 24-well plate (Greiner Bio-One). The initial concentrations of each of the strains wt_agg_, wt_agg2_*ΔpilE*, wt_plank_, wt_plank2_ were approximately 3 × 10^6^ cells ml^-1^. Cells were grown for 10 h at 37°C, 5% C0_2_ in an Infinite M200 plate reader with a shaking period of 2 min per OD cycle (OD was measured every 10 min). Microscopy samples were taken to assess aggregation of each culture at this time point. Batch effects were minimized by using three replicates of each culture from the same day.

Total RNA was isolated from liquid gonococcal cultures with the RNeasy Mini Kit (Qiagen) according to the manufacturer’s instructions. Prior to RNA isolation, the bacterial cultures were mixed with the RNA protect Bacteria Reagent (Qiagen) according to the manufacturer’s instructions and the pellets were stored at −80 °C.

Next-generation sequencing, in particular the Illumina HiSeq system (NEB), was used to obtain gonococcal transcriptomes. mRNA sequencing was carried out by the Cologne Center for Genomics (University of Cologne, Germany) with 100 bp paired-end reads and on average 10 million reads per sample. For sample submission, the final RNA concentration was adjusted to 50-200 ng/μl in at least 10 μl volume.

### Raw read analysis pipeline and differential gene expression

As a reference, the MS11 genome sequence (Ngo MS11) was obtained from NCBI with the accession number CP003909.1. This was used as a reference for all strains in the fasta format and the annotation was downloaded in the gff3 format. For improving the MS11 annotation, the orthologs of all genes were detected in *N. gonorrhoeae* FA1090 (Ngo FA1090) with the accession number NC_002946.2 and *N. meningitidis* MC85 (Nmen) NC_003112.2 via blastn (64).

The sequencing reads from each data set were trimmed and paired using Trimmomatic (version 0.36) (65). The reads were then mapped against the reference genome of Neisseria gonorrhoeae MS11 (NCBI, CP003909.1) using STAR (2.5.3a) (66). Subsequently, read counts were called for each gene of the reference with featureCounts from the subread package (version: 2.0.1) (67). The package DESeq2 implemented in R was used to perform the principal component analysis with the plotPCA function and raw count data was transformed with the regularized logarithm function. We calculated the differential gene expression with the DESeq2 package implemented in R. For each gene, we obtained the log2-fold change of the biofilm-forming strain wt_agg_ with respect to each planktonic strain ΔpilE, wt_pilE17_, wt_pilE24_, wt_pilE32_ as well as the associated adjusted *p*-value. A gene is regarded as differentially regulated if the (adjusted) *p*-value is ≤ 0.05 and the | log2-fold change| ≥ 0.5.

Genes with redundant regions across the genome were excluded from the analysis. These genes were detected by blasting single gene sequences against the Ngo MS11 genome and detecting regions where blast coverage was at least at 30% with an identity of 99% or higher. For Ngo MS11, we find 130 multi-mapping genes, which are excluded from the further analysis. As the *pilS* loci and *pilE* share similar regions among each other, we excluded all of these loci from the analysis of the global transcriptome.

### Functional annotation with KEGG categories

As Ngo MS11 is not functionally annotated well, we use the Ngo FA1090 strain and its KEGG annotation (38). Each gene is sorted into one of 17 categories. Large categories containing more than 60 genes include: phage associated, metabolism of co-factors and vitamins, hypothetical, genetic information processing, energy metabolism, cellular processes/ human disease, carbohydrate metabolism, and amino acid metabolism. Genes collected in the category ’hypothetical’ are not assumed to share any biological function. In the enrichment analysis, the genes that have no orthologue in Ngo FA1090 or that are not connected to any gene category in Ngo FA1090 are excluded. We test for the significance of functional enrichment in each category by performing a one-sided Fisher’s exact test and Bonferroni correction to mitigate the multiple testing problem. Categories were condensed such that each gene is associated with one category.

### Manuscript editing

DeepL was used for editing the grammar and formulations of the manuscript.

## Acknowledgements

We thank Gerrit Ansmann, Tobias Bollenbach, Natalie Balaban, and the Maier group for helpful discussions. This work has been supported by the Center for Molecular Medicine Cologne, the Deutsche Forschungsgemeinschaft through grant MA3898 and CRC1310, and the IHRS BioSoft.

## Supplementary Figures

**S1 Fig.:**
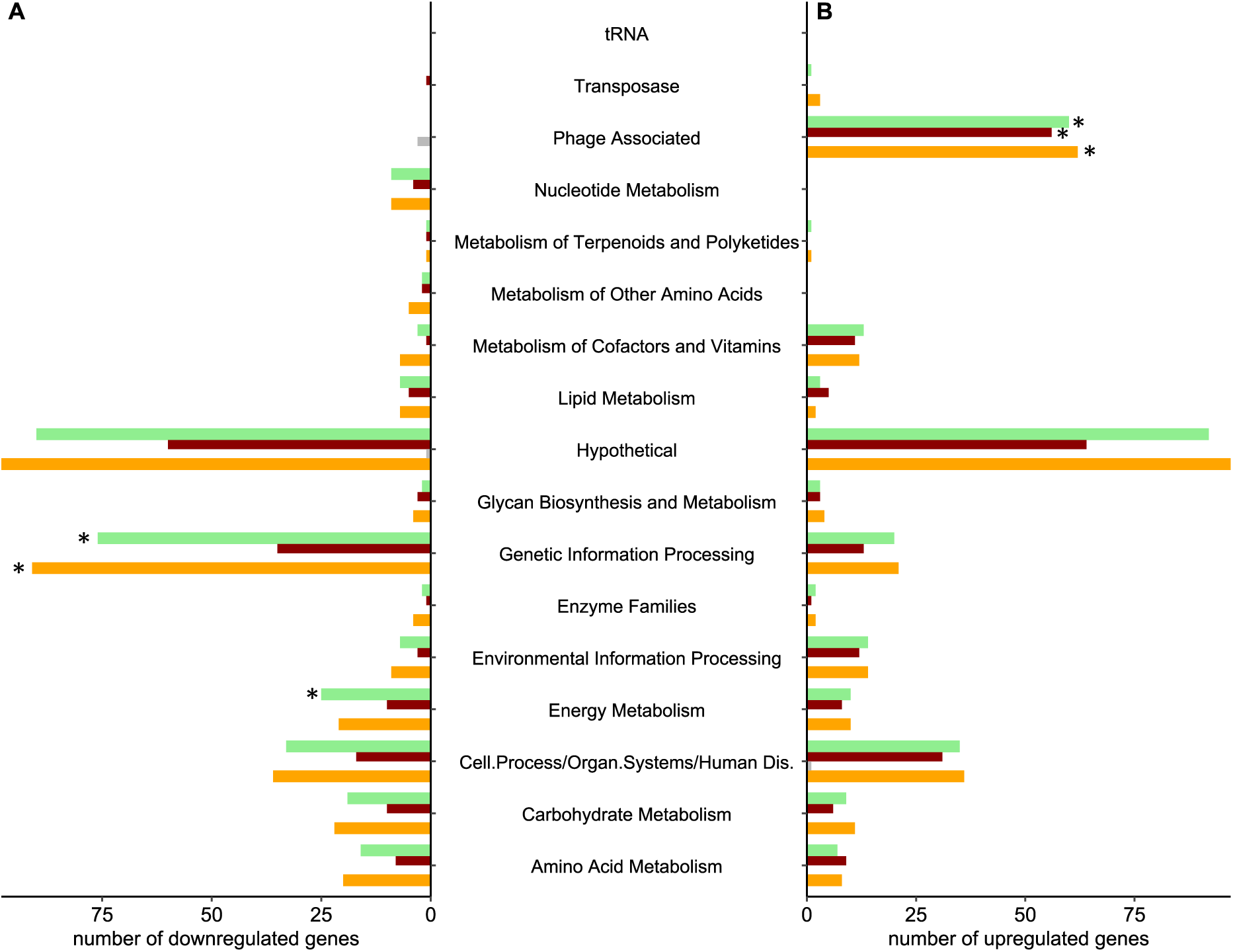
Functional enrichment of differentially expressed genes, wt_agg_ vs. wt_plank_ (orange), wt_plank2_ (red), and *ΔpilE* (green). Genes that belonged to neither KEGG category nor had an FA1090 ortholog are excluded. Only significantly differentially expressed genes are shown. A) Number of downregulated genes. B) Number of upregulated genes. Significantly enriched categories are highlighted with * (p-value ≤ 0.01).

**S2 Fig.:**
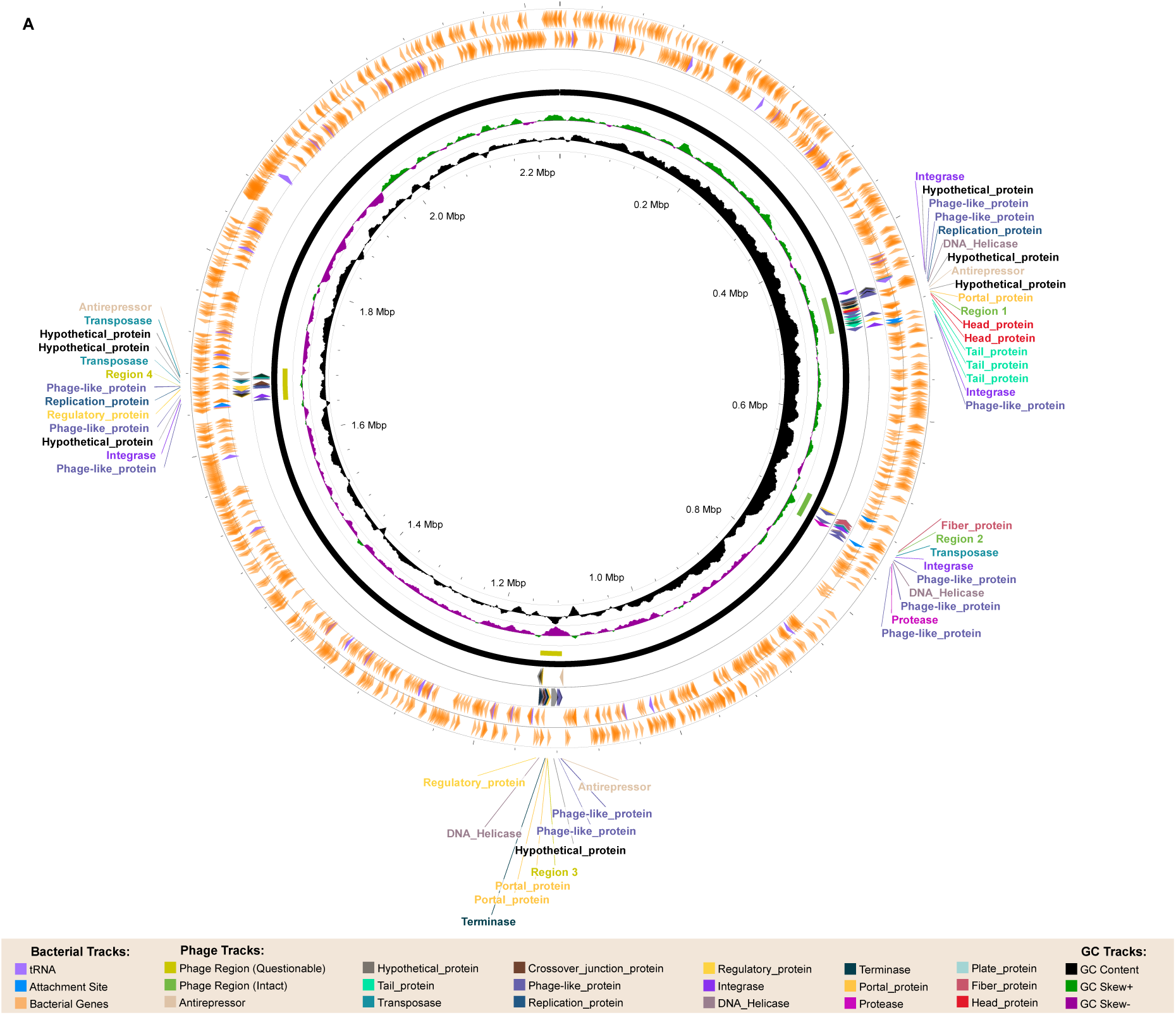
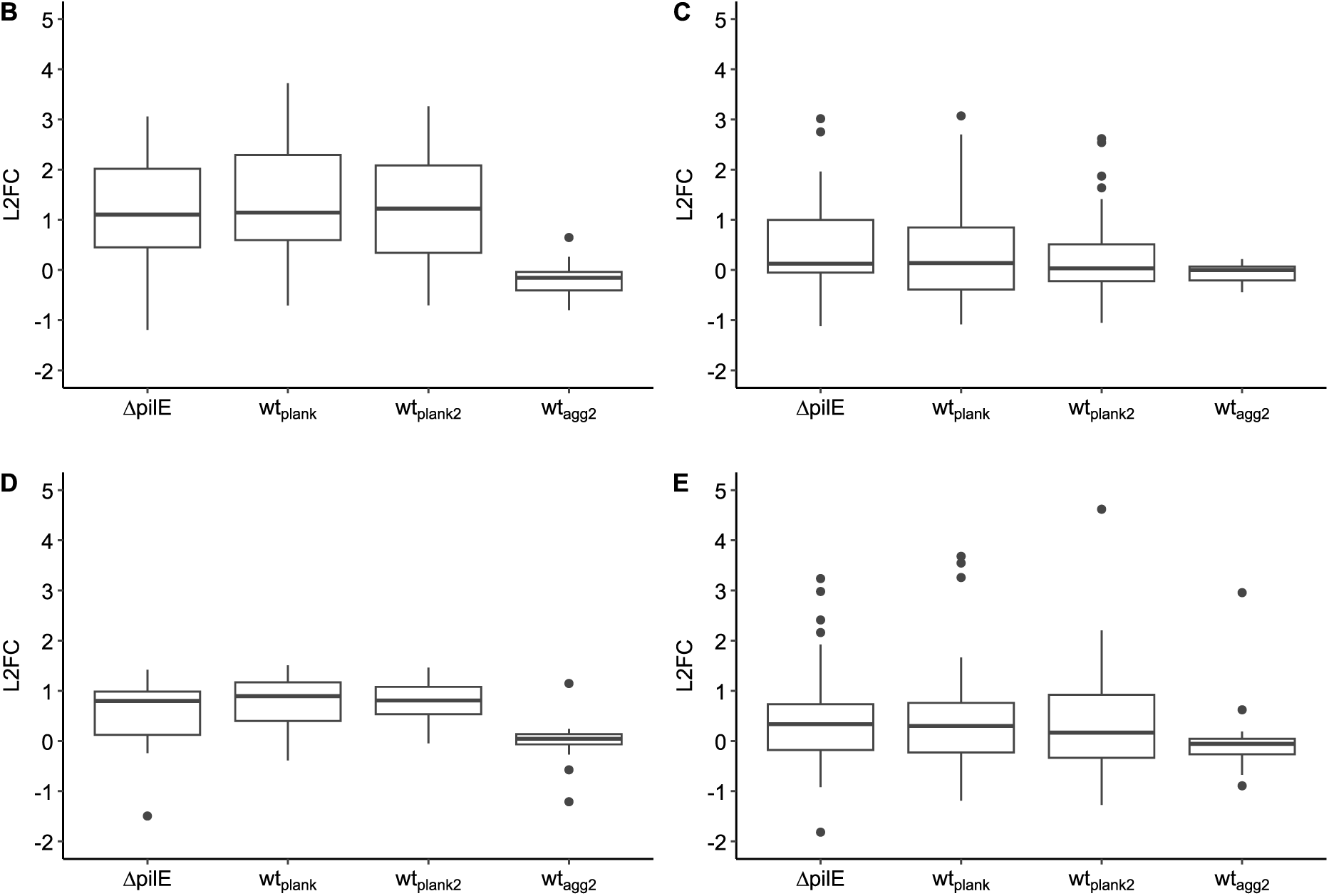
Prophages of *N. gonorrhoeae* MS11 (CP003909.1) as predicted with PHASTEST (39) and their differential expression. L2FC were obtained by comparing the expression of *pilE* variant strains to wt_agg_. Genes were categorized according to the four predicted prophages of *N. gonorrhoeae* MS11 by PHASTEST. All genes between the respective genome range were assigned as part of the corresponding phage. A) Circular map of the *N. gonorrhoeae* MS11 (CP003909.1) genome with positions of predicted prophages. Differential expression of B) Φ1, C) Φ2, D) Φ3, and E) Φ4.

**S3 Fig.:**
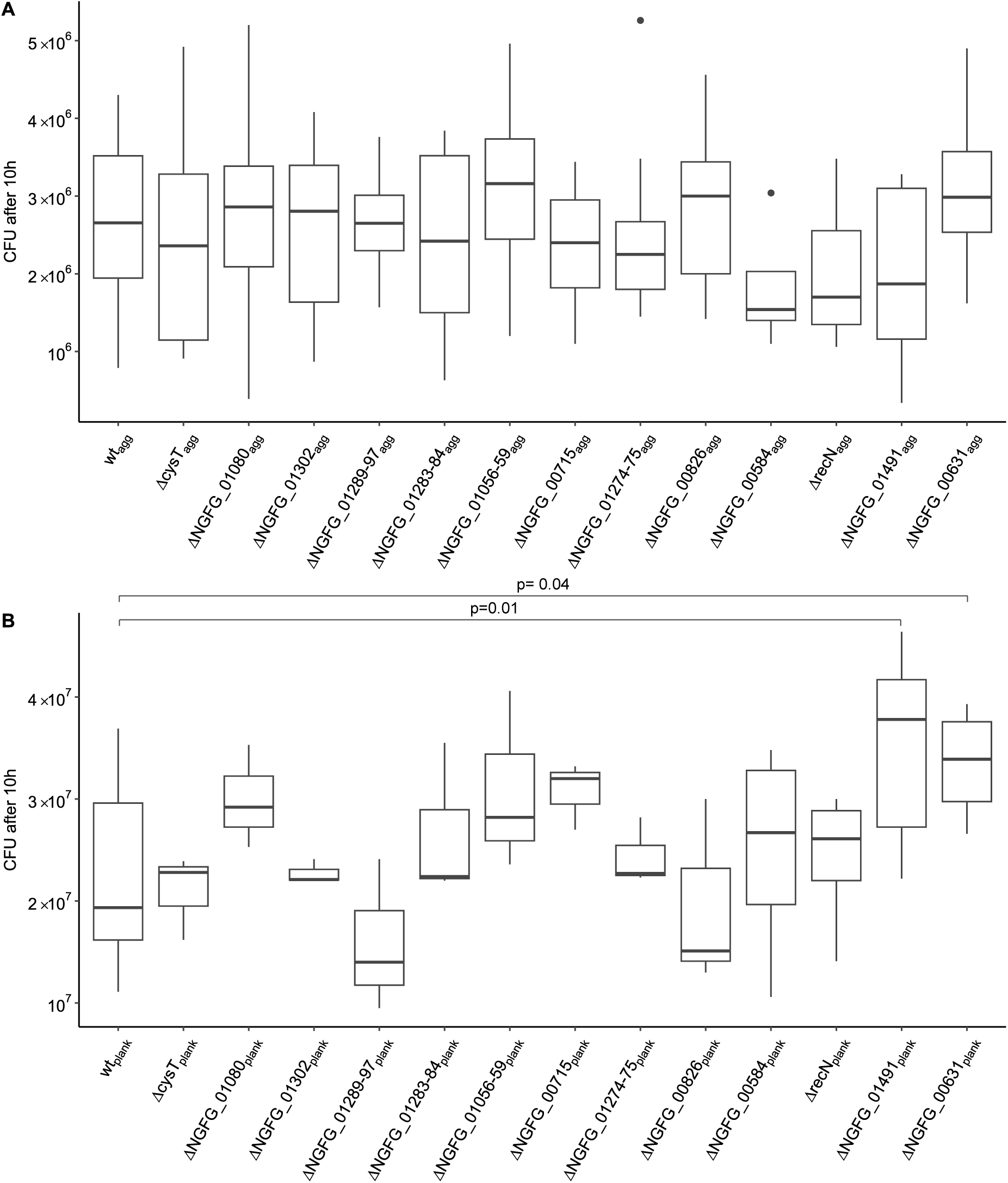
CFU of A) aggregating and B) planktonic strains after 10 h of growth in liquid.

**S4 Fig.:**
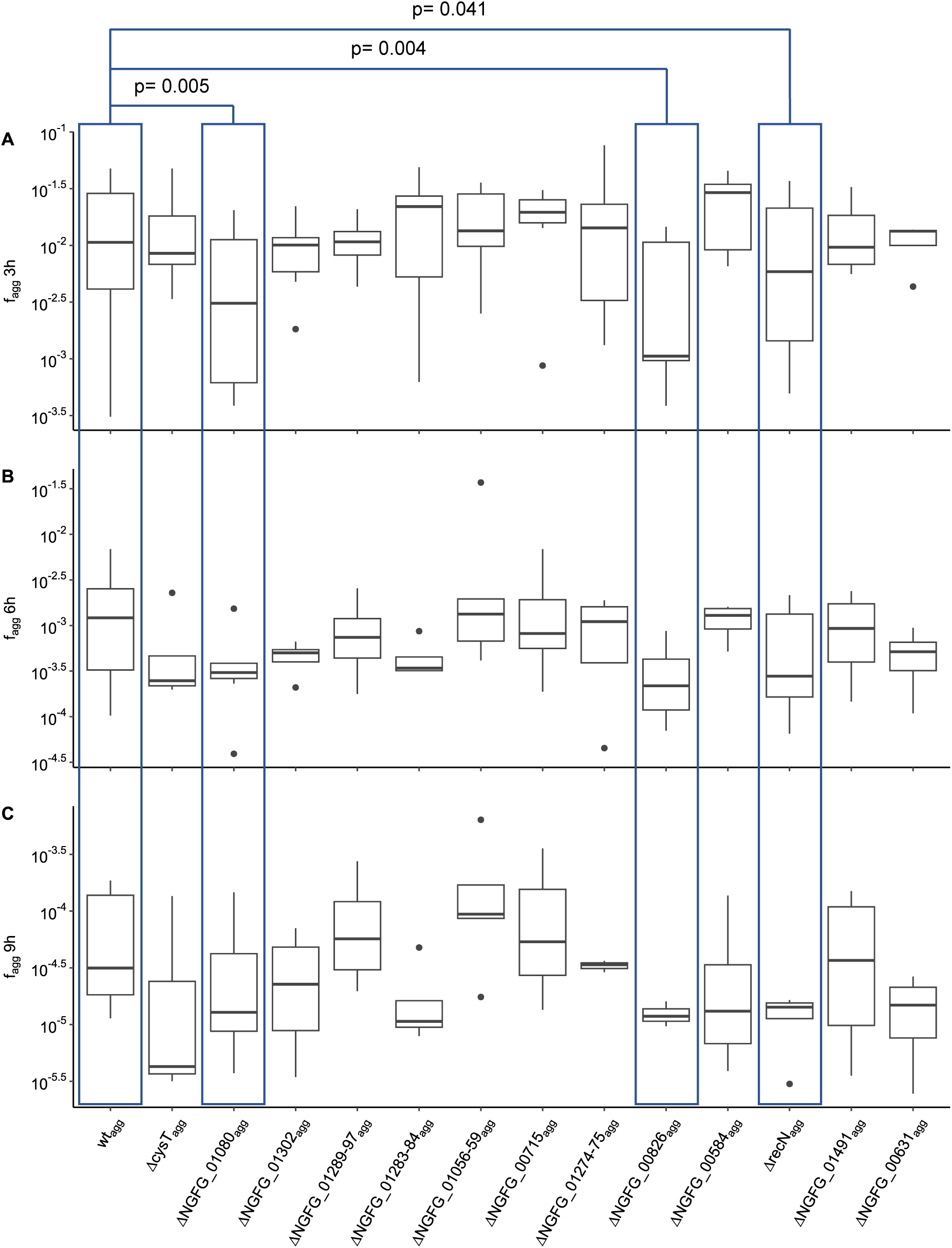
Fractions of surviving aggregating cells under ciprofloxacin treatment for A) 3 h, B) 6 h and C) 9 h.

**S5 Fig.:**
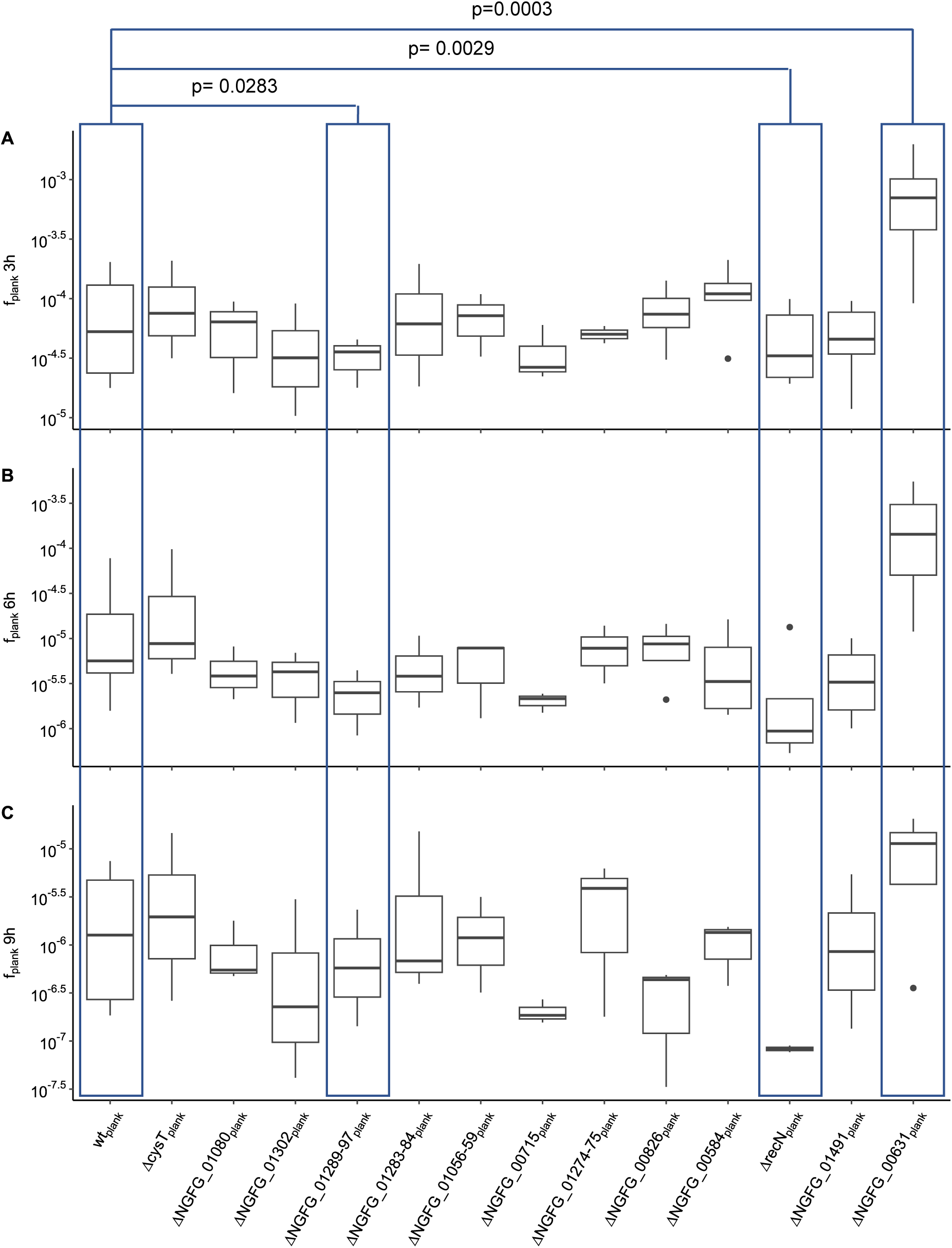
Fractions of surviving planktonic strains under ciprofloxacin treatment for A) 3 h, B) 6 h and C) 9 h.

**S6 Fig.:**
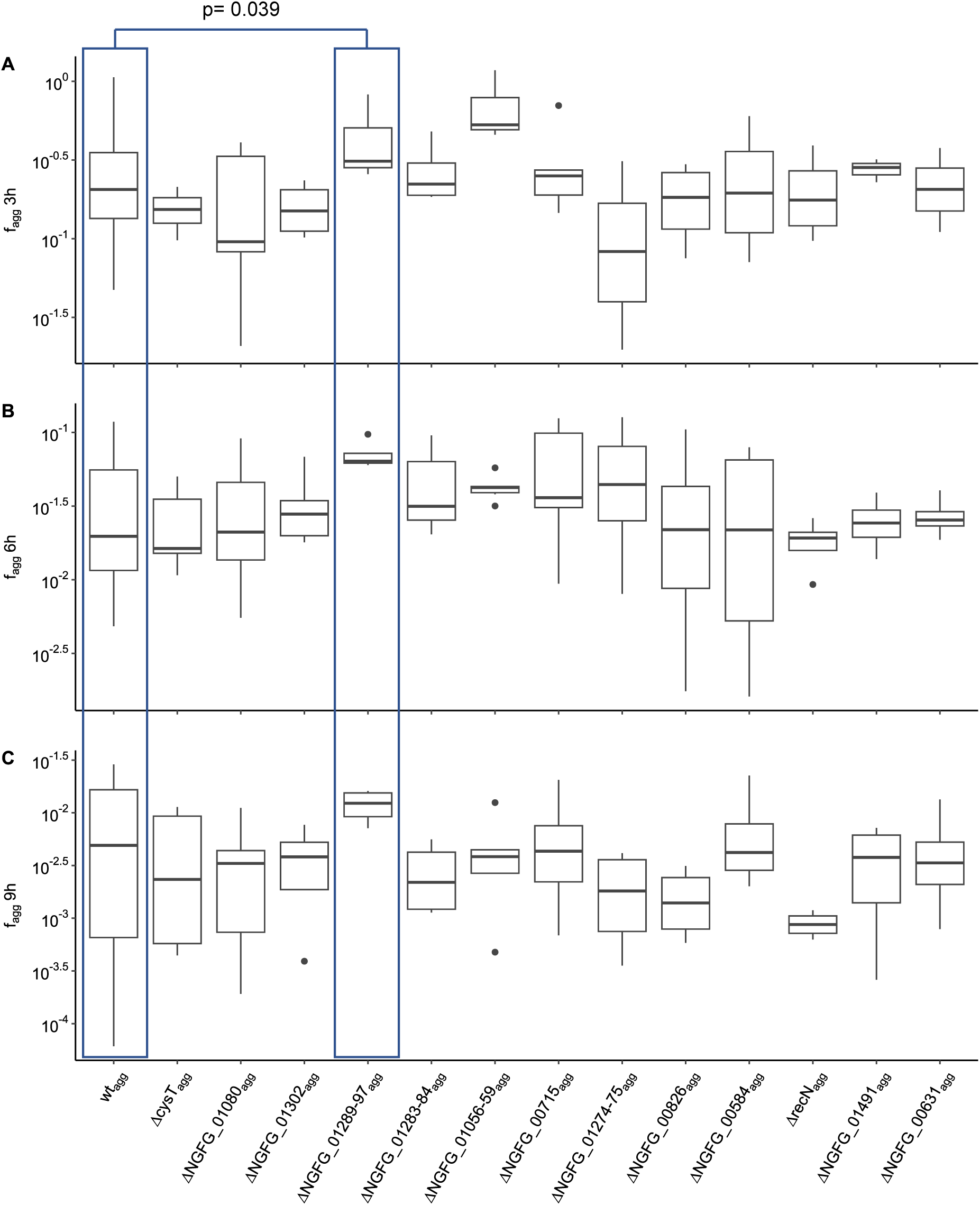
Fractions of surviving aggregating cells under ceftriaxone treatment for A) 3 h, B) 6 h and C) 9 h.

**S7 Fig.:**
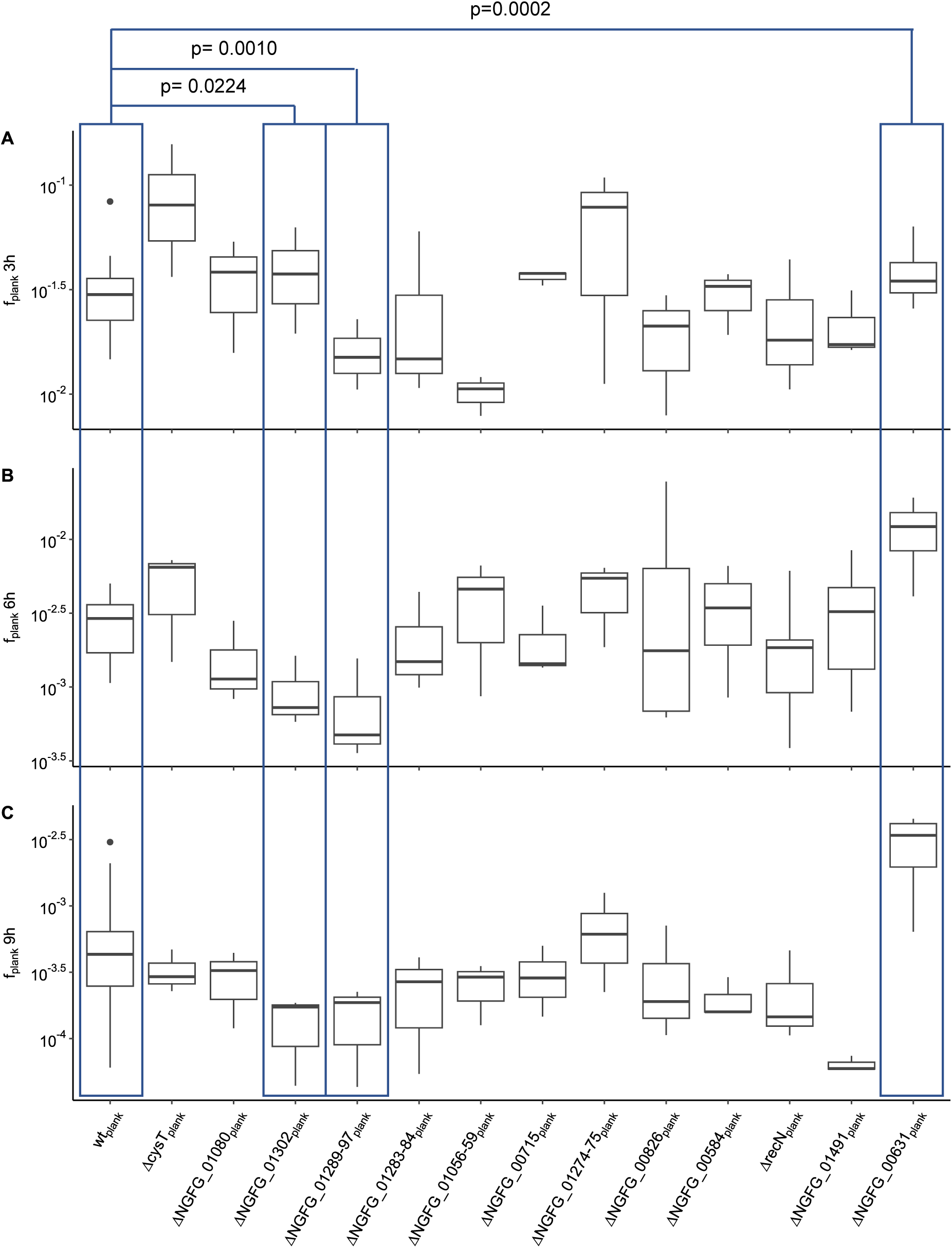
Fractions of surviving planktonic cells under ceftriaxone treatment for A) 3 h, B) 6 h and C) 9 h.

**S8 Fig.:**
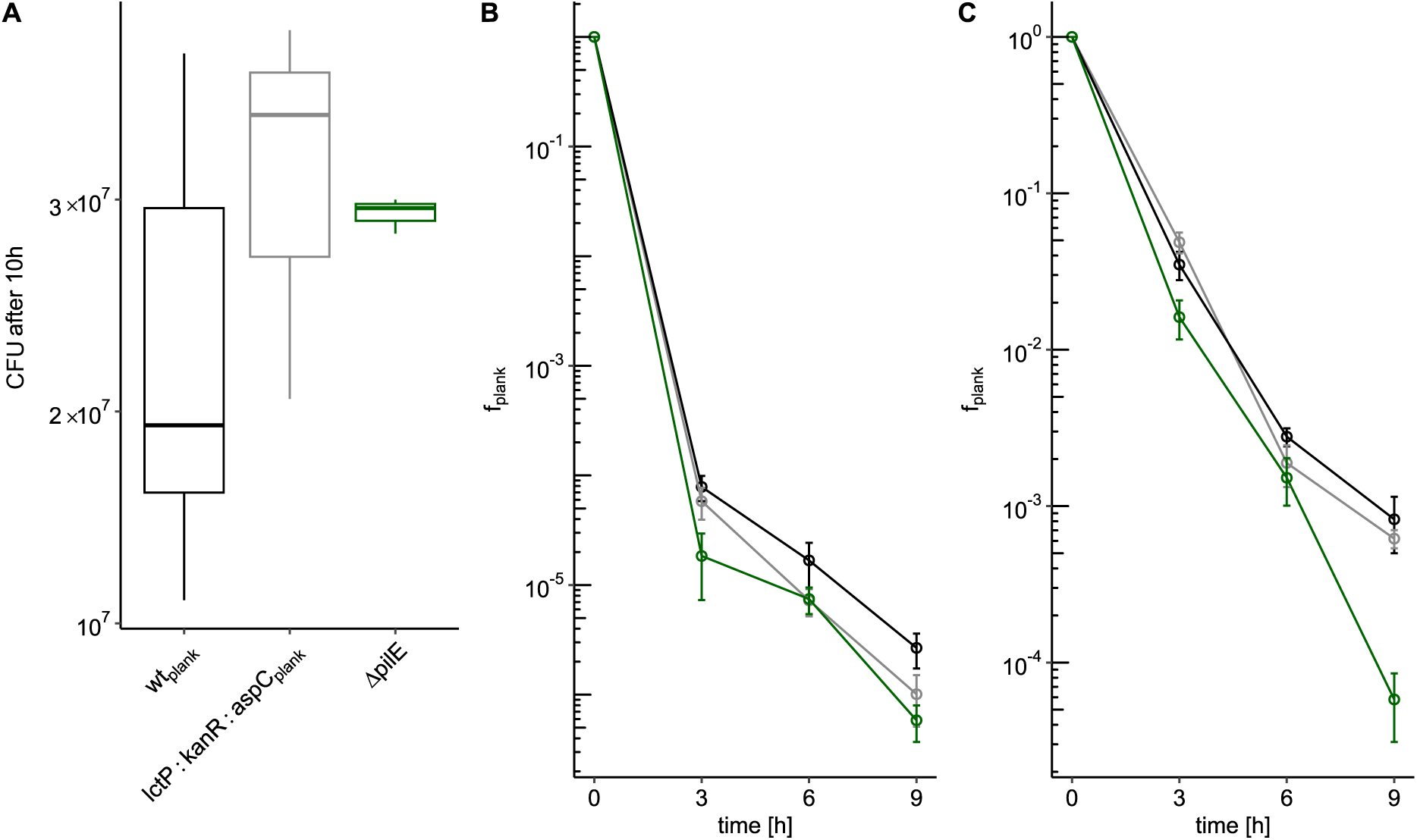
The impact of *pilE* deletion and the kanamycin resistance cassette on cell survival during ciprofloxacin and ceftriaxone treatment compared to wt_plank_. *kanR* was introduced into the intergenic *lctP::aspC* locus. A) CFU after 10 h of strains wt_plank_ (Ng240, black), *ΔpilE* (Ng253, green), lctP:kanR:aspC_plank_ (Ng290, grey). B, C) Fractions of surviving planktonic control strains and their respective parental strain. wt_plank_ (Ng240, black), *ΔpilE* (Ng253, green), lctP:kanR:aspC_plank_ (Ng290, grey) under B) ciprofloxacin (p_wtplank-ΔpilE_= 0.12, p_wtplank-lctP:kanR:aspCplank_= 0.87) and C) ceftriaxone treatment (p_wtplank-ΔpilE_= 9.8×10^-4^, p_wtplank-lctP:kanR:aspCplank_= 0.76).

**S9 Fig.:**
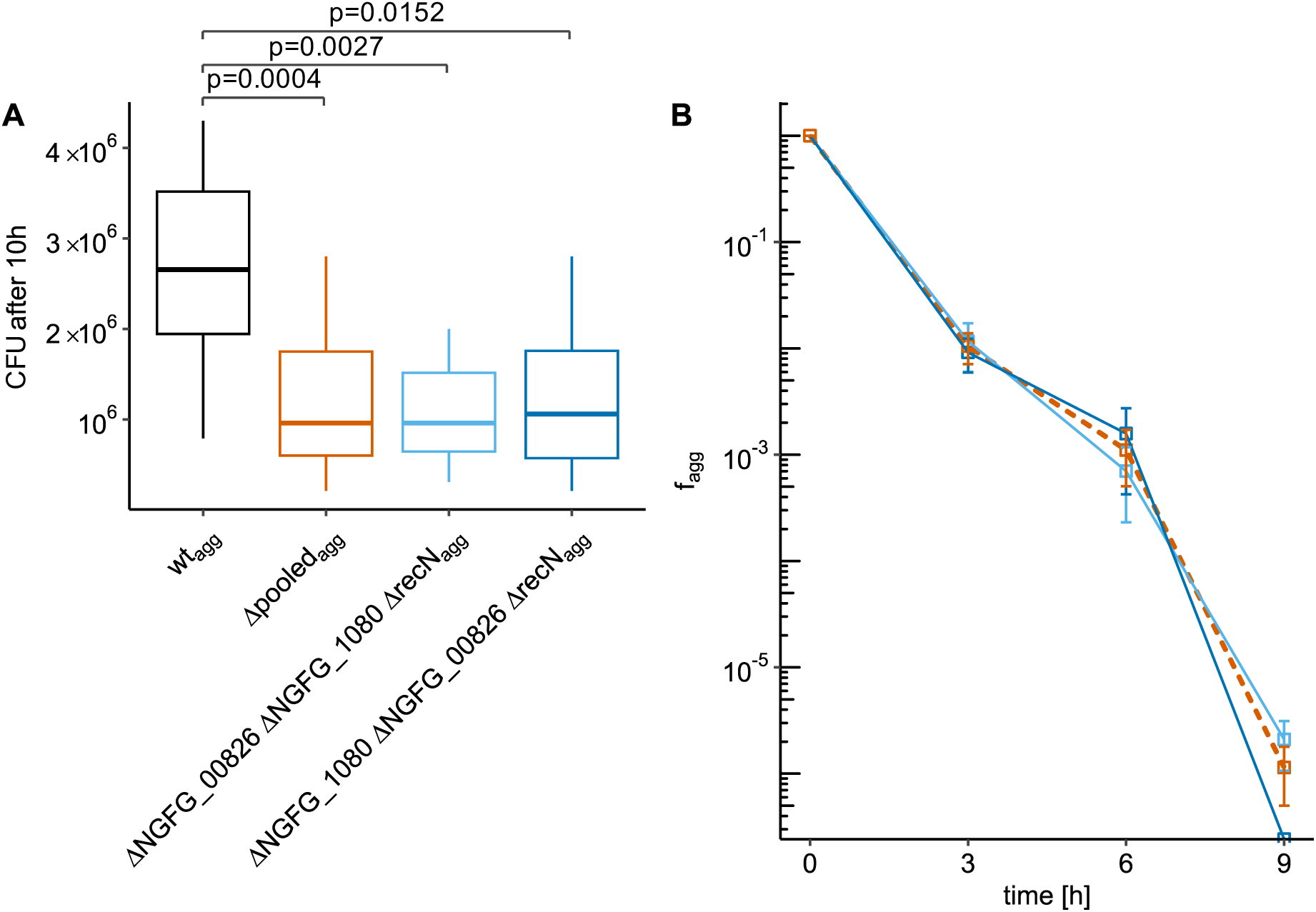
Triple knock out strains. The order of the deletion does not significantly impact the triple knock out phenotype. Therefore, we pooled the data of both genotypes for better statistics. A) CFU after 10 h. B) Fractions of surviving aggregating triple KO strains ΔNGFG_00826 ΔNGFG_01080 Δ*recN*_agg_ (Ng321, cyan) and ΔNGFG_01080 ΔNGFG_00826 Δ*recN*_agg_ (Ng325, blue) compared to the pooled data (dashed, orange). p-value between ΔNGFG_01080 Δ*recN*_agg_ (Ng321) and ΔNGFG_01080 ΔNGFG_00826 Δ*recN*_agg_ (Ng325) = 0.14.

**S10 Fig.:**
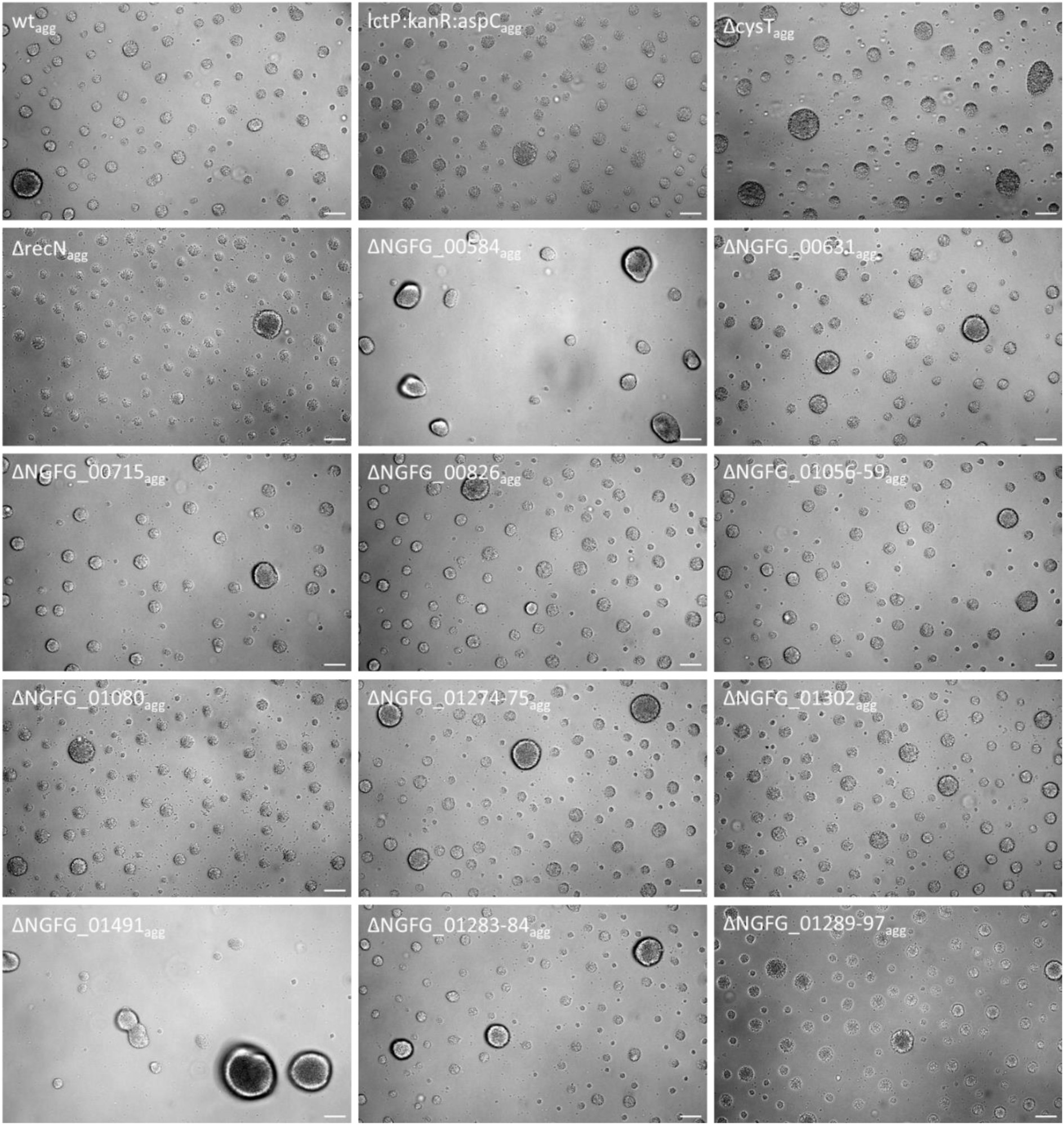
All mutants in the wt_agg_ background maintain the ability to form aggregates in liquid media. Cells were grown for 5 h without shaking (37°C, 5% CO_2_). Scale bar: 25 µm.

## Supplementary tables and data sheets

**S1 Table:**
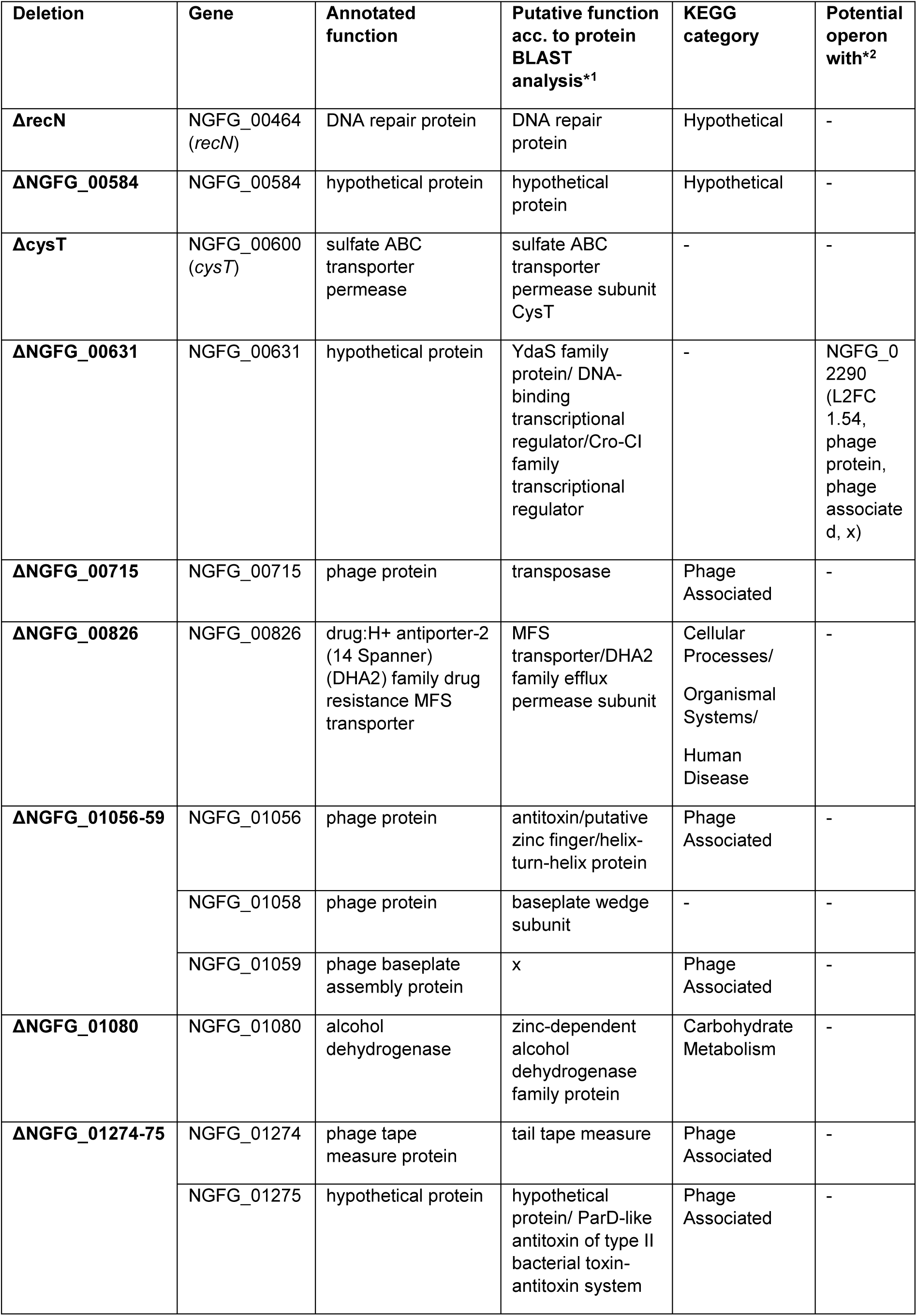
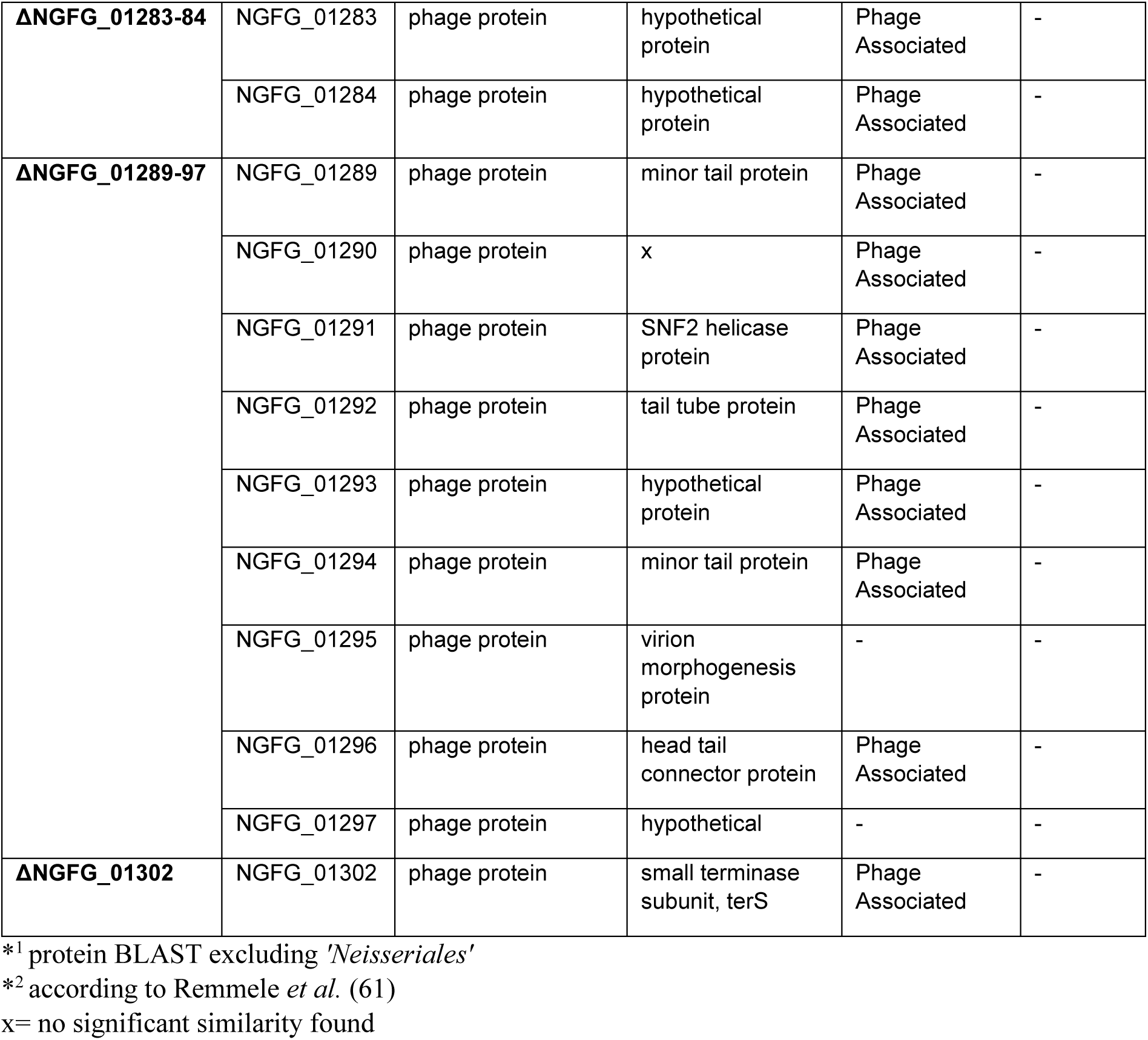
Overview of gene deletion strains.

| Deletion | Gene | Annotated function | Putative function acc. to protein BLAST analysis*1 | KEGG category | Potential operon with*2 |
| --- | --- | --- | --- | --- | --- |
| **ΔrecN** | NGFG_00464 (*recN*) | DNA repair protein | DNA repair protein | Hypothetical | - |
| **ΔNGFG_00584** | NGFG_00584 | hypothetical protein | hypothetical protein | Hypothetical | - |
| **ΔcysT** | NGFG_00600 (*cysT*) | sulfate ABC transporter permease | sulfate ABC transporter permease subunit CysT | - | - |
| **ΔNGFG_00631** | NGFG_00631 | hypothetical protein | YdaS family protein/ DNA-binding transcriptional regulator/Cro-CI family transcriptional regulator | - | NGFG_02290 (L2FC 1.54, phage protein, phage associated, x) |
| **ΔNGFG_00715** | NGFG_00715 | phage protein | transposase | Phage Associated | - |
| **ΔNGFG_00826** | NGFG_00826 | drug:H<sup>+</sup> antiporter-2 (14 Spanner) (DHA2) family drug resistance MFS transporter | MFS transporter/DHA2 family efflux permease subunit | Cellular Processes/<br>Organismal Systems/<br>Human Disease | - |
| **ΔNGFG_01056-59** | NGFG_01056 | phage protein | antitoxin/putative zinc finger/helix-turn-helix protein | Phage Associated | - |
|  | NGFG_01058 | phage protein | baseplate wedge subunit | - | - |
|  | NGFG_01059 | phage baseplate assembly protein | x | Phage Associated | - |
| **ΔNGFG_01080** | NGFG_01080 | alcohol dehydrogenase | zinc-dependent alcohol dehydrogenase family protein | Carbohydrate Metabolism | - |
| **ΔNGFG_01274-75** | NGFG_01274 | phage tape measure protein | tail tape measure | Phage Associated | - |
|  | NGFG_01275 | hypothetical protein | hypothetical protein/ ParD-like antitoxin of type II bacterial toxin-antitoxin system | Phage Associated | - |
| **ΔNGFG_01283-84** | NGFG_01283 | phage protein | hypothetical protein | Phage Associated | - |
|  | NGFG_01284 | phage protein | hypothetical protein | Phage Associated | - |
| **ΔNGFG_01289-97** | NGFG_01289 | phage protein | minor tail protein | Phage Associated | - |
|  | NGFG_01290 | phage protein | x | Phage Associated | - |
|  | NGFG_01291 | phage protein | SNF2 helicase protein | Phage Associated | - |
|  | NGFG_01292 | phage protein | tail tube protein | Phage Associated | - |
|  | NGFG_01293 | phage protein | hypothetical protein | Phage Associated | - |
|  | NGFG_01294 | phage protein | minor tail protein | Phage Associated | - |
|  | NGFG_01295 | phage protein | virion morphogenesis protein | - | - |
|  | NGFG_01296 | phage protein | head tail connector protein | Phage Associated | - |
|  | NGFG_01297 | phage protein | hypothetical | - | - |
| **ΔNGFG_01302** | NGFG_01302 | phage protein | small terminase subunit, terS | Phage Associated | - |

\*<sup>1</sup> protein BLAST excluding '*Neisseriales*'
\*<sup>2</sup> according to Remmele *et al.* (61)
x= no significant similarity found

**S2 Table:** MIC of aggregating strains.

| Strain | MIC [$\mu\text{g/ml}$] | |
| --- | --- | --- |
|  | ceftriaxone | ciprofloxacin |
| **wt<sub>agg</sub>** | 0.008 | 0.004 |
| **$\Delta\text{cysT}_{\text{agg}}$** | 0.008 | 0.004 |
| **$\Delta\text{NGFG\_01080}_{\text{agg}}$** | 0.008 | 0.004 |
| **$\Delta\text{NGFG\_01302}_{\text{agg}}$** | 0.008 | 0.004 |
| **$\Delta\text{NGFG\_01289-97}_{\text{agg}}$** | 0.008 | 0.004 |
| **$\Delta\text{NGFG\_01283-84}_{\text{agg}}$** | 0.008 | 0.004 |
| **$\Delta\text{NGFG\_01056-59}_{\text{agg}}$** | 0.008 | 0.004 |
| **$\Delta\text{NGFG\_00715}_{\text{agg}}$** | 0.008 | 0.004 |
| **$\Delta\text{NGFG\_01274-75}_{\text{agg}}$** | 0.008 | 0.004 |
| **$\Delta\text{NGFG\_00826}_{\text{agg}}$** | 0.008 | 0.004 |
| **$\Delta\text{NGFG\_00584}_{\text{agg}}$** | 0.008 | 0.004 |
| **$\Delta\text{recN}_{\text{agg}}$** | 0.008 | 0.002 |
| **$\Delta\text{NGFG\_01491}_{\text{agg}}$** | 0.008 | 0.004 |
| **$\Delta\text{NGFG\_00631}_{\text{agg}}$** | 0.008 | 0.004 |
| **lctP:kanR:aspC<sub>agg</sub>** | 0.008 | 0.004 |
| **3x$\Delta_{\text{agg}}$** | not determined | 0.004 |

**S3 Table:** Strains used in this study.

| Abbreviation | Strain number | Strain | Relevant genotype | Reference |
| --- | --- | --- | --- | --- |
| Wt<sub>agg</sub> | Ng150 | ΔG4 (wt*) | *G4::apraR* | (29) |
| - | Ng225 | T126C step1 | *G4::apraR*,<br>*pilE::pilE<sup>T126C</sup>* *ermC*<br>*rpsL<sub>s</sub>* | (27) |
| - | Ng052 | Δ*comE*<sub>234</sub>Δ*pilV* | *comE4::Kan*,<br>*comE3::CIm*,<br>*comE2::Erm*,<br>*recA6ind(tetM) pilVfs* | (68) |
| Wt<sub>plank2</sub> | Ng230 | Wt<sub>pilE32</sub> | *G4::apraR*,<br>*pilE<sub>clinicalisolateNG32</sub>* | (37) |
| Wt<sub>plank</sub> | Ng240 | Wt<sub>pilE17</sub> | *G4::apraR*,<br>*pilE<sub>clinicalisolateNG17</sub>* | (37) |
| Wt<sub>agg2</sub> | Ng242 | Wt<sub>pilE24</sub> | *G4::apraR*,<br>*pilE<sub>clinicalisolateNG24</sub>* | (37) |
| Δ*pilE* | Ng253 | Δ*pilE* | *pilE::kanR* | (37) |
| ΔrecN<sub>agg</sub> | Ng261 | ΔG4 ΔrecN | *recN::kanR* | this study |
| ΔNGFG_00826<sub>agg</sub> | Ng262 | ΔG4 ΔNGFG_00826 | *NGFG_00826::kanR* | this study |
| ΔNGFG_01491<sub>agg</sub> | Ng263 | ΔG4 ΔNGFG_01491 | *NGFG_01491::kanR* | this study |
| ΔNGFG_00584<sub>agg</sub> | Ng264 | ΔG4 ΔNGFG_00584 | *NGFG_00584::kanR* | this study |
| ΔNGFG_01283-84<sub>agg</sub> | Ng269 | ΔG4 ΔNGFG_01283-1284 | *NGFG_01283-1284::kanR* | this study |
| ΔNGFG_01274-1275<sub>agg</sub> | Ng270 | ΔG4 ΔNGFG_01274-1275 | *NGFG_01274-1275::kanR* | this study |
| ΔNGFG_01056-1059<sub>agg</sub> | Ng271 | ΔG4 ΔNGFG_01056-1059 | *NGFG_01056-1059::kanR* | this study |
| ΔNGFG_00715<sub>agg</sub> | Ng272 | ΔG4 ΔNGFG_00715 | *NGFG_00715::kanR* | this study |
| ΔNGFG_01289-97<sub>agg</sub> | Ng273 | ΔG4 ΔNGFG_01289-1297 | *NGFG_01289-1297::kanR* | this study |
| ΔrecN<sub>plank</sub> | Ng275 | NG17 ΔrecN | *recN::kanR* | this study |
| ΔNGFG_00826<sub>plank</sub> | Ng276 | NG17 ΔNGFG_00826 | *NGFG_00826::kanR* | this study |
| ΔNGFG_01491<sub>plank</sub> | Ng277 | NG17 ΔNGFG_01491 | *NGFG_01491::kanR* | this study |
| ΔNGFG_00584<sub>plank</sub> | Ng278 | NG17 ΔNGFG_00584 | *NGFG_00584::kanR* | this study |
| ΔcysT<sub>agg</sub> | Ng279 | ΔG4 ΔcysT | *cysT::kanR* | this study |
| ΔNGFG_01302<sub>agg</sub> | Ng280 | ΔG4 ΔNGFG_01302 | *NGFG_01302::kanR* | this study |
| ΔNGFG_01080<sub>agg</sub> | Ng281 | ΔG4 ΔNGFG_01080 | *NGFG_01080::kanR* | this study |
| ΔNGFG_00631<sub>plank</sub> | Ng288 | NG17 ΔNGFG_00631 | *NGFG_00631::kanR* | this study |
| ΔNGFG_00631<sub>agg</sub> | Ng289 | ΔG4 ΔNGFG_00631 | *NGFG_00631::kanR* | this study |
| *lctP::kanR::aspC*<sub>plank</sub> | Ng290 | NG17 *lctP::kanR::aspC* | *lctP::kanR::aspC* | this study |
| *lctP::kanR::aspC*<sub>agg</sub> | Ng291 | ΔG4 *lctP::kanR::aspC* | *lctP::kanR::aspC* | this study |
| ΔNGFG_01284-83<sub>plank</sub> | Ng296 | NG17 ΔNGFG_01284-1283 | *NGFG_01284-1283::kanR* | this study |
| ΔNGFG_01274-75<sub>plank</sub> | Ng297 | NG17 ΔNGFG_01274-1275 | *NGFG_01274-1275::kanR* | this study |
| ΔNGFG_01056-59<sub>plank</sub> | Ng298 | NG17 ΔNGFG_01056-1059 | *NGFG_01056-1059::kanR* | this study |
| ΔNGFG_00715<sub>plank</sub> | Ng299 | NG17 ΔNGFG_00715 | *NGFG_00715::kanR* | this study |
| ΔNGFG_01289-97<sub>plank</sub> | Ng300 | NG17 ΔNGFG_01289-1297 | *NGFG_01289-1297::kanR* | this study |
| ΔcysT<sub>plank</sub> | Ng301 | NG17 ΔcysT | *cysT::kanR* | this study |
| ΔNGFG_01302<sub>plank</sub> | Ng302 | NG17 ΔNGFG_01302 | *NGFG_01302::kanR* | this study |
| ΔNGFG_01080<sub>plank</sub> | Ng303 | NG17 ΔNGFG_01080 | *NGFG_01080::kanR* | this study |
| ΔNGFG_00826<br>ΔNGFG_01080<br>ΔrecN<sub>agg</sub> | Ng321 | ΔG4 ΔNGFG_00826<br>ΔNGFG_01080 ΔrecN | *NGFG_00826::kanR*<br>*ΔNGFG_01080 ΔrecN* | this study |
| ΔNGFG_01080<br>ΔNGFG_00826<br>ΔrecN<sub>agg</sub> | Ng325 | ΔG4 ΔNGFG_01080<br>ΔNGFG_00826 ΔrecN | *NGFG_01080::kanR*<br>*ΔNGFG_00826 ΔrecN* | this study |

**S4 Table:** Primer combinations used for *kanR* replacements.

| Gene | Primer A | Primer B | Primer C | Primer D | Primer E | Primer F |
| --- | --- | --- | --- | --- | --- | --- |
| NGFG_00464 | sk260 | sk261 | sk262 | sk263 | sk264 | sk265 |
| NGFG_00826 | sk248 | sk249 | sk250 | sk251 | sk252 | sk253 |
| NGFG_01491 | sk266 | sk267 | sk268 | sk269 | sk270 | sk271 |
| NGFG_00584 | sk254 | sk255 | sk256 | sk257 | sk258 | sk259 |
| NGFG_01284, 1283 | sk288 | sk289 | sw28 | sw29 | sk290 | sk291 |
| NGFG_01275, 1274 | sk292 | sk293 | sw28 | sw29 | sk294 | sk295 |
| NGFG_01059, 1058, 1056 | sk296 | sk297 | sw28 | sw29 | sk298 | sk299 |
| NGFG_00715 | sk304 | sk305 | sw28 | sw29 | sk306 | sk307 |
| NGFG_01289-1297 | sk308 | sk309 | sw28 | sw29 | sk310 | sk311 |
| NGFG_00600 | sk344 | sk345 | sw28 | sw29 | sk346 | sk347 |
| NGFG_01302 | sk340 | sk341 | sw28 | sw29 | sk342 | sk343 |
| NGFG_01080 | sk348 | sk349 | sw28 | sw29 | sk350 | sk351 |
| NGFG_00631 | sk370 | sk371 | sw28 | sw29 | sk372 | sk373 |
| lctP:aspC | tc14 | sk381 | sw28 | sw29 | sk382 | sk383 |

**S5 Table:**
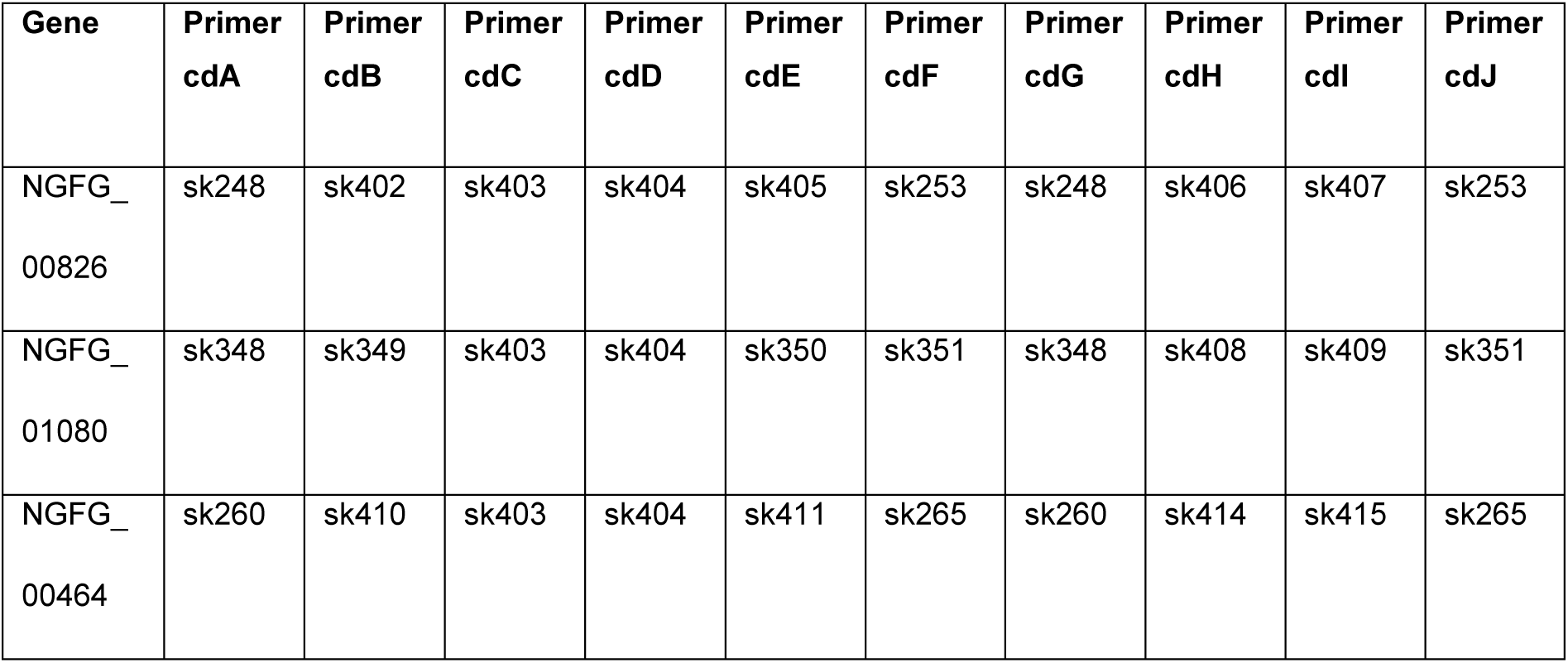
Primer combinations used for clean deletions.

**S6 Table:**
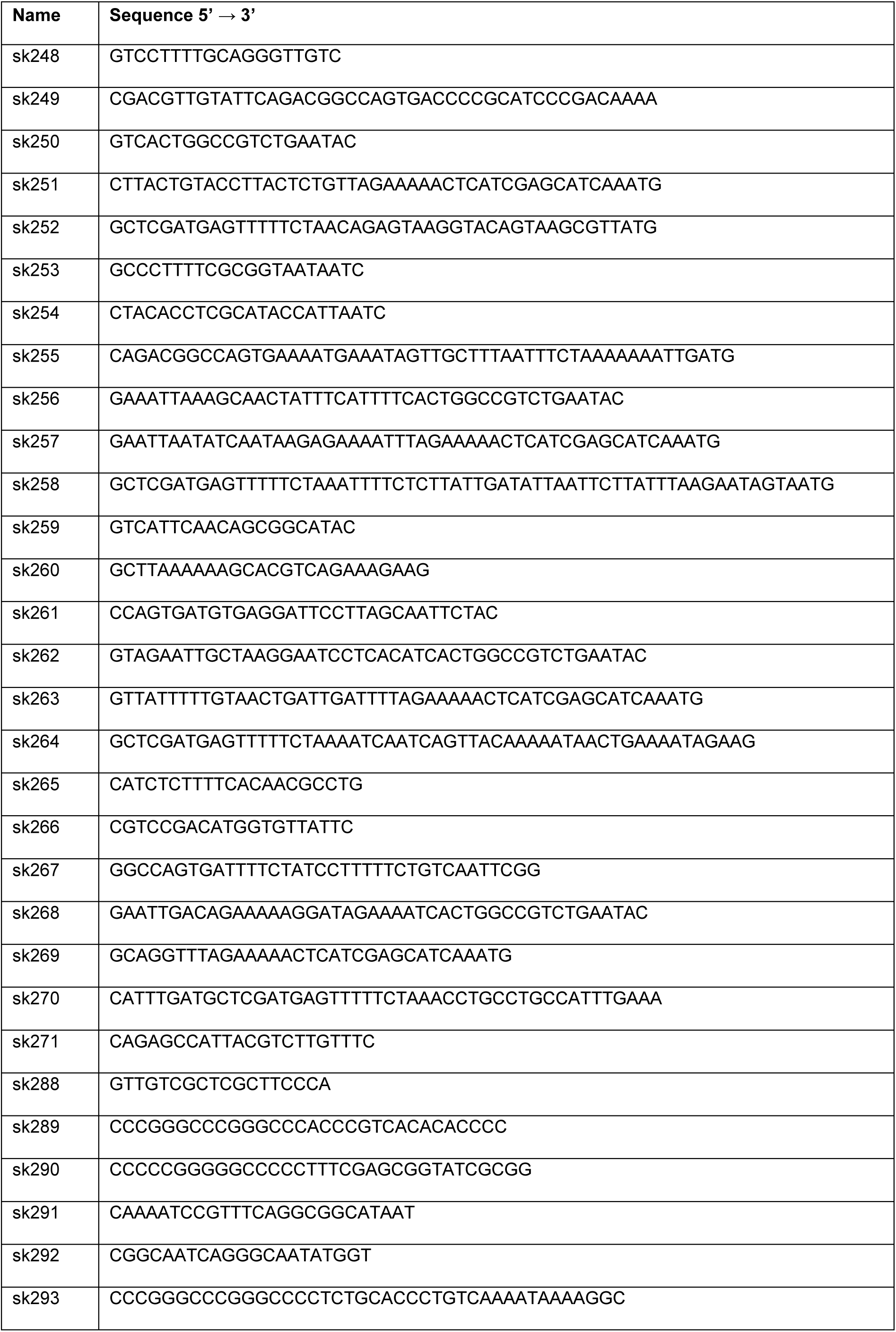
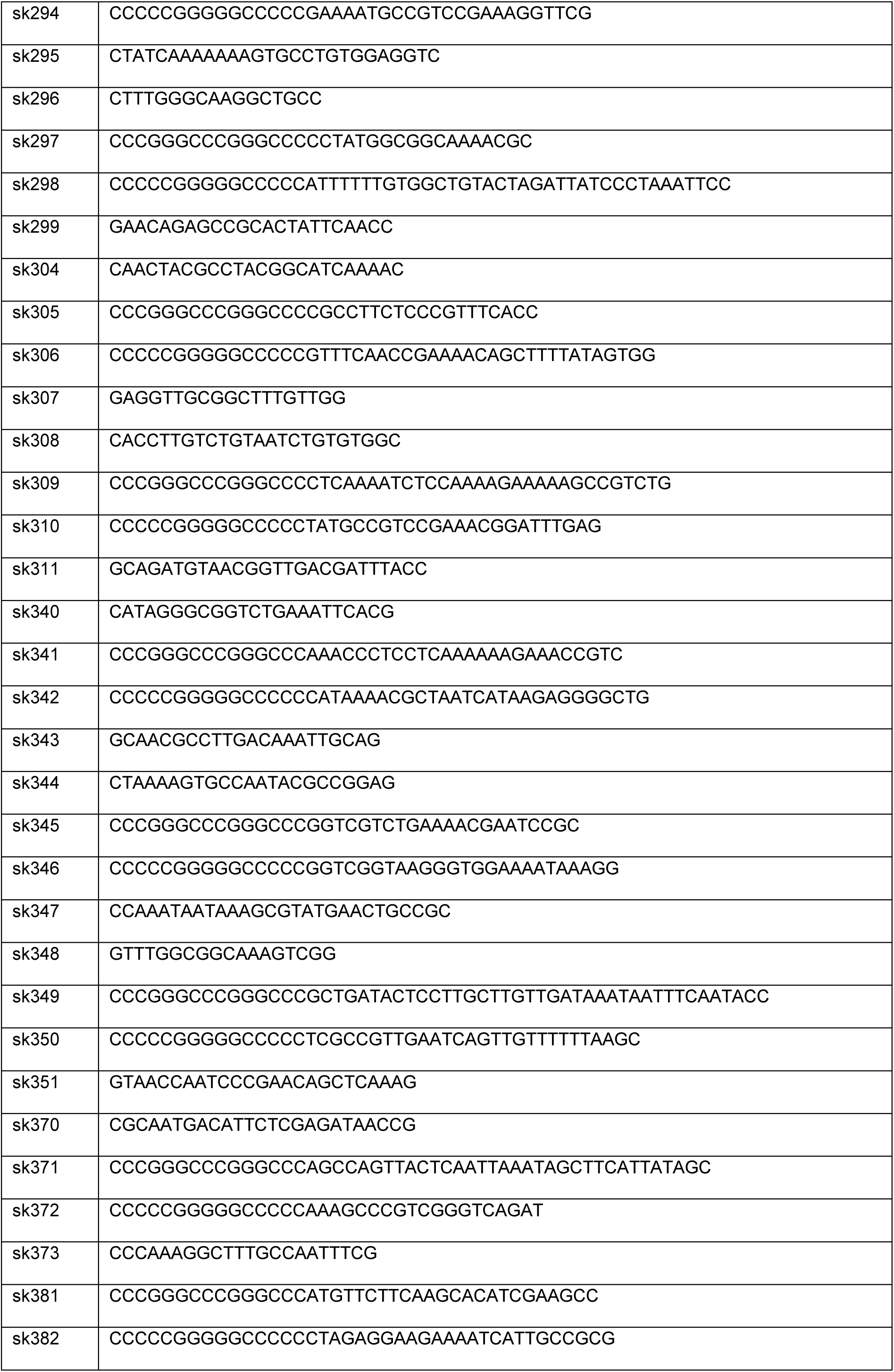
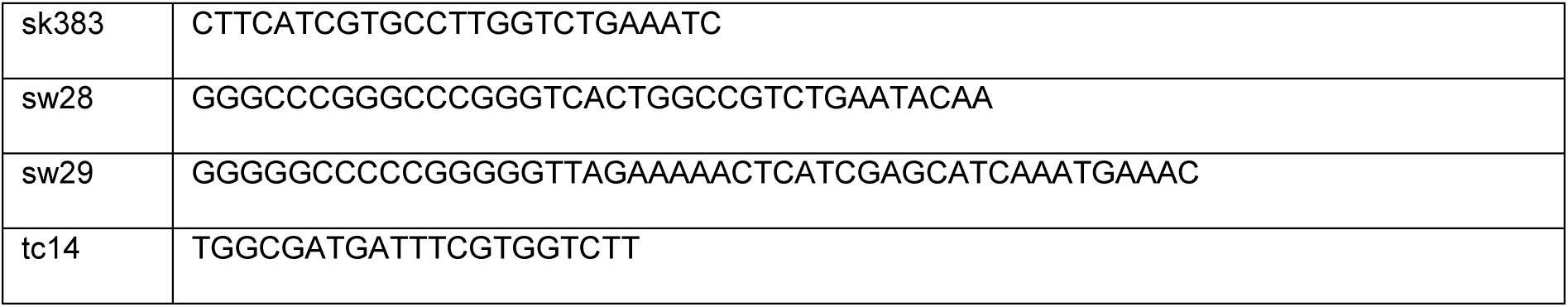
Primers used in this study.

